# Aberrant miR-29 is a predictive feature of severe phenotypes in pediatric Crohn’s disease

**DOI:** 10.1101/2022.12.16.520635

**Authors:** Alexandria J. Shumway, Michael T. Shanahan, Emilie Hollville, Kevin Chen, Caroline Beasley, Jonathan W. Villanueva, Sara Albert, Grace Lian, Moises R. Cure, Matthew Schaner, Lee-Ching Zhu, Surekha Bantumilli, Mohanish Deshmukh, Terrence S. Furey, Shehzad Z. Sheikh, Praveen Sethupathy

## Abstract

Crohn’s disease (CD) is a chronic inflammatory gut disorder. Molecular mechanisms underlying the clinical heterogeneity of CD remain poorly understood. MicroRNAs (miRNAs) are important regulators of gut physiology and several have been implicated in the pathogenesis of adult CD. However, there is a dearth of large-scale miRNA studies for pediatric CD. We hypothesized that specific miRNAs uniquely mark pediatric CD. We performed small RNA-sequencing of patient-matched non-inflamed colon and ileum biopsies from treatment-naïve pediatric patients with CD (n=169) and a control cohort (n=108). Comprehensive miRNA analysis revealed 58 miRNAs altered in pediatric CD. Notably, multinomial logistic regression analysis revealed that index levels of ileal miR-29 are strongly predictive of severe inflammation and stricturing. Transcriptomic analyses of transgenic mice overexpressing miR-29 show a significant reduction of the tight junction protein gene *Pmp22* and classic Paneth cell markers. The dramatic loss of Paneth cells was confirmed by histologic assays. Moreover, we also found that pediatric CD patients with elevated miR-29 exhibit significantly lower Paneth cell counts, increased inflammation scores, and reduced levels of *PMP22*. These findings strongly indicate that miR-29 up-regulation is a distinguishing feature of pediatric CD, highly predictive of severe phenotypes, and associated with inflammation and Paneth cell loss.

## Introduction

Crohn’s disease (CD) represents one of the primary inflammatory bowel diseases (IBD) and is thought to develop due to dysregulated inflammatory responses in genetically susceptible individuals. Over the course of the past decade especially, CD has become an increasingly global disease, with growing incidence in newly industrialized countries ^1–3^. The number of CD cases is predicted to increase in the US by almost 1.5-fold by 2025^4^. Some reports have attributed this growth in substantive part to the increase in CD cases among the pediatric population, which has more than doubled since the start of the 21^st^ century and remains the fastest-growing affected age group ^5–8^.

The precise causes of CD are still enigmatic, but it is believed to be an aberrant immune response to a complex interaction of factors including environmental exposures, genetics, and the gut microbiome. Both adult and pediatric CD are characterized by non-contiguous lesions in the gastrointestinal tract that can lead to chronic abdominal pain, diarrhea, fistulas, and/or abscesses. Up to 30% of CD patients are pediatric, and these patients tend to exhibit a more severe phenotype due to concomitant issues of growth failure, poor bone density, and delayed puberty^9,10^. Although for some patients the existing therapies can aid in mucosal healing to decrease the need for surgical intervention and improve the overall quality of life, there is currently no cure ^10,11^. Treatment of CD can differ greatly across patients since the appropriate therapeutic regimen relies on many factors including location and behavior of disease, comorbidities, previous treatments, and age ^12,13^. Remission of CD remains difficult to achieve, especially in pediatric patients^12^. The complexity and heterogeneity of CD and the highly variable efficacy of existing therapeutic options highlight the need for novel intervention methods. The development of novel biomarkers and prognostic indicators could further aid clinicians in determining disease trajectory and response to therapy^14^.

In recent years, we and others have investigated microRNAs (miRNAs) as potential diagnostic markers, prognostic indicators of disease severity, and candidate therapeutic targets for adult CD^15–17^. MiRNAs are small non-coding RNAs (∼22 nts long) that post-transcriptionally regulate gene expression and have been shown to influence most major biological processes and diseases^18^. These molecules regulate the majority of protein-coding genes and each miRNA can target up to hundreds of mRNAs, resulting in mRNA destabilization or inhibition of translation^18,19^. Dysfunction of miRNA activity can lead to acute or chronic inflammation, which is characteristic of CD ^20,21^. In 2010, a seminal study showed the contribution of overall miRNA activity to intestinal architecture and function ^20^. More recently, specific miRNAs have been implicated in IBD ^22,23,24^. For example, we showed that a single miRNA (miR-31) is a major driver of the differences between two molecular subtypes of adult CD^25^. Further functional studies demonstrated that miR-31 regulates barrier function in part by controlling the expression of activin A receptor-like type 1 (ALK1) and that high levels of miR-31 are strongly associated with poor clinical outcomes and increased likelihood of relapse after remission^26^. This study, as well as several others, have established miRNAs as valuable prognostic indicators of disease behavior and potential therapeutic targets in adult CD^15,24,27^.

Despite these advances, a major limitation of these studies is that they pertain largely to adult CD. There are only a handful of studies that focus on miRNAs in pediatric CD, and most of these do not use a sequencing-based approach to define comprehensive miRNA profiles ^24,27–33^. Recently, we performed the first large-scale sequencing of miRNAs in pediatric CD, but we focused the analysis on only one miRNA^25^. Here, we substantially expand the cohort, and perform a comprehensive quantitative analysis of all miRNAs in matched ileum and colon samples from pediatric patients and non-IBD controls. We discover and report on specific miRNAs that exhibit the greatest utility as predictive molecular features of key clinical outcomes in pediatric CD. We also perform functional follow-up studies in vivo of one particular miRNA that is a distinguishing feature of pediatric CD (relative to adult CD) and define new potential targets and functions that merit further investigation.

## Results

### Ileal and colonic microRNA profiles stratify pediatric CD from non-IBD patients

We had previously performed small RNA-sequencing (smRNA-seq) on 60 ileum and 76 colon tissue index biopsies from pediatric CD patients as well as 50 ileum and 48 colon tissue samples from non-IBD (NIBD) individuals ^25^. In that study, we focused our analysis on only one miRNA of interest. Therefore, the full potential of the predictive power of miRNAs for clinical outcomes in pediatric CD remained unknown. To fill this important knowledge gap, and to define a comprehensive miRNA signature of pediatric CD, we first expanded the cohort to 277 total samples and then implemented and applied the bioinformatic analysis pipeline miRquant 2.0. Subsequent to the implementation of this pipeline, we removed datasets with less than one million reads mapped to miRNAs (Methods) (Figure 1A, Table 1). We also removed from further analysis those samples from individuals without detailed clinical information, including SIM, age, and sex (Methods). The age and sex breakdown of the pediatric patients whose samples remained for further analysis are provided in Table 2 (n=245).

**Table 1.** Sample breakdown with thresholds applied.

|  | Colon CD | Ileum CD | Colon NIBD | Ileum NIBD |
| --- | --- | --- | --- | --- |
| Original | 93 | 76 | 54 | 54 |
| Pass 1 million reads mapped to miRs | 87 | 74 | 53 | 53 |
| Have clinical data | 75 | 65 | 52 | 53 |

**Table 2.** Demographics of pediatric CD patients.

| Patient Characteristics | Colon CD (Average) | Ileum CD (Average) | Colon NIBD (Average) | Ileum NIBD (Average) |
| --- | --- | --- | --- | --- |
| Diagnosis Age (in years) | 12 | 11.85 | 12.27 | 12.34 |
| Grouped Ages |  |  |  |  |
| VEO (<6) | 2 (2.7%) | 3 (4.6%) | 8 (15.4%) | 8 (15.1%) |
| Child (7-12) | 36 (48%) | 31 (47.6%) | 17 (32.7%) | 17 (32.1%) |
| Teen (13-17) | 37 (49.3%) | 31 (47.6%) | 27 (51.9%) | 28 (52.8%) |
| Sex |  |  |  |  |
| Female | 27 (36%) | 22 (33.8%) | 31 (59.6%) | 32 (60.4%) |
| Male | 48 (64%) | 43 (66.2%) | 21 (40.4%) | 21 (39.6%) |

**Figure 1.**
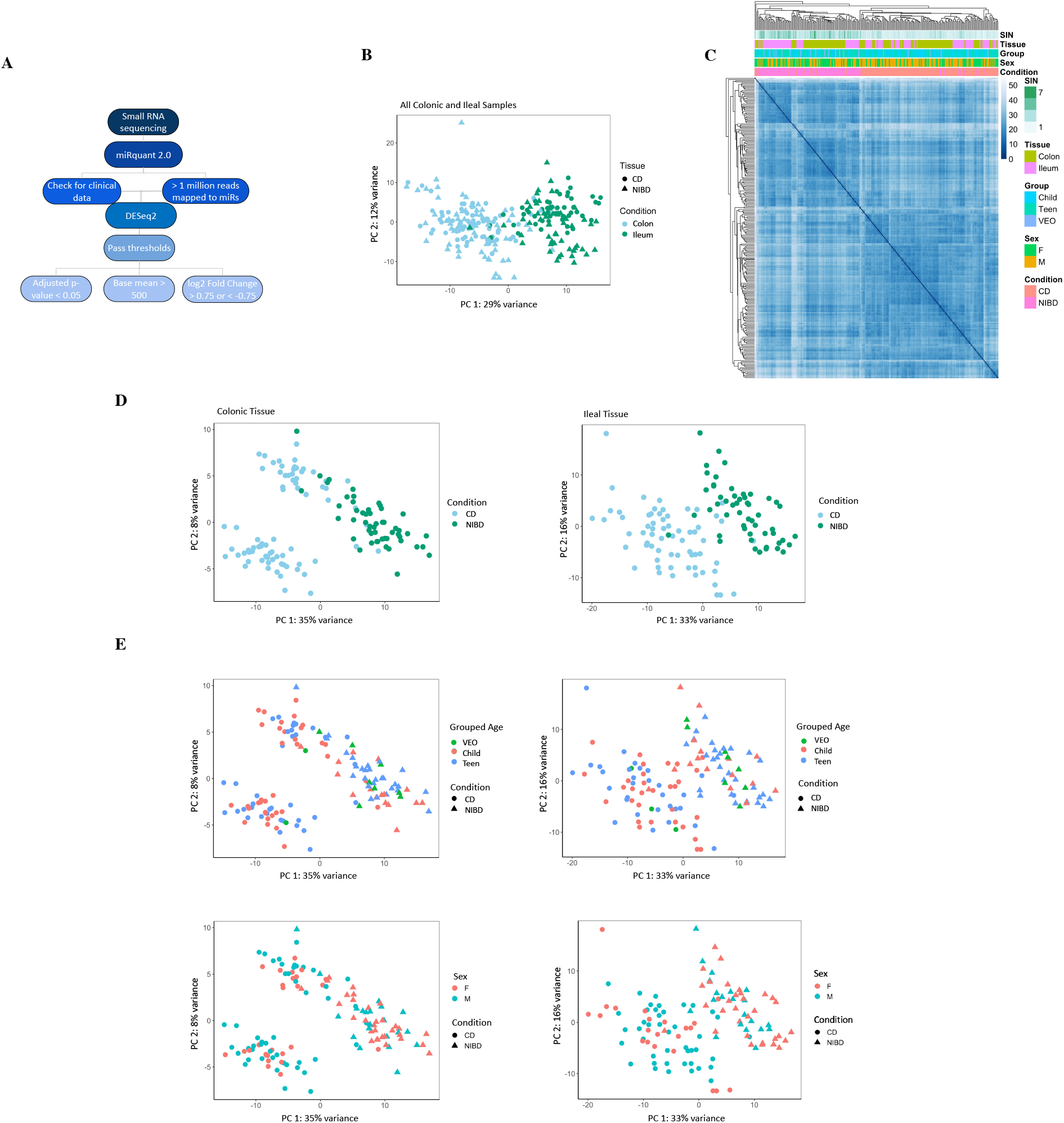
Ileal and colonic microRNA profiles stratify by disease status. (A) Workflow for small-RNA sequencing analysis. (B) Principal component analysis (PCA) of VST normalized counts for all (n=245) CD and NIBD samples accounting for the covariates of small-RNA integrity metric (SIM), grouped ages (VEO < 6, Child = 7-12, Teen = 13-17), and sex. The CD samples are represented as circles and NIBD samples as squares. The two tissue types are colored in blue and green for colon and ileum, respectively. The percent of variation explained is indicated for principal component 1 along the x-axis and principal component 2 along the y-axis. (C) Unsupervised hierarchical clustering of the Euclidean distances among pediatric samples (n=245) was calculated based on VST normalized counts. The miRNA profiles for the colon (left) and ileal tissue (right) account for the covariates of SIM, grouped ages, and sex. The CD and NIBD samples are indicated by peach and pink boxes. Other covariates are represented as the colors indicated by the legend. (D) PCA of pediatric miRNA profiles in colon (n=127) (left) and ileal (n=118) (right) tissue accounting for the covariates of SIM, grouped ages, and sex. (E) PCA plots for colon (left) and ileum (right) in which the grouped patient age and sex phenotype information are overlaid. The colors green, red, and blue represent the VEO, Child, and Teen group ages, respectively. Female and male patients are indicated by the color red and blue, respectively. Disease status is specified by shape.

Unsupervised hierarchical clustering and principal component analysis (PCA) of the miRNA profiles across all 245 samples showed a strong stratification by disease status (CD vs. NIBD) (Figure 1B-C). This result was maintained when considering only the patient-matched samples (n=228) (Figure S1). PCA of the colonic and ileal miRNA datasets, separately, also revealed a robust grouping of samples by disease (CD vs. NIBD) along the first principal component (Figure 1D). There was no additional stratification detected according to age or sex (Figure 1E).

### Colonic microRNAs separate pediatric CD into two clusters

Performing PCA analysis on each tissue individually, pediatric CD patients appear to be stratified into two clusters based on colonic miRNA profiles (Figure 1D), while this is not clearly evident with ileal miRNA profiles (Figure 1D). Intriguingly, it is apparent that the clustering among colonic CD cases is partially due to future ileal stricturing status (Figure S2A). To define the colonic miRNAs that may explain this stratification, we sought to identify the miRNAs that are significantly differentially expressed between the two clusters (Figure S2B). Using DESeq2 we found two colonic miRNAs (miR-99b and miR-146b) significantly more highly expressed in cluster #1, which includes all but one of the individuals who developed ileal stricturing, and one colonic miRNA (miR-451a) significantly more highly expressed in cluster #2 (Figure S2C).

### Many but not all microRNAs significantly altered in pediatric CD are shared between ileum and colon

Our data affords the unique opportunity to define the miRNAs that are enriched in either the human colon or the ileum at baseline, using matched tissue from the same control (NIBD) individuals. Using DESeq2 we found 10 miRNAs significantly differentially abundant between the ileum and colon (Figure 2A) – 7 miRNAs significantly enriched in the ileum (Figure 2B) and 3 miRNAs significantly enriched in the colon (Figure 2C). The ileal-enriched miRNAs include miR-31, which we have previously reported on, most prominently in adult Crohn’s^25,26^. The colon-enriched miRNAs include miR-196b, which has long been associated with colonic CD ^23,34^ and ulcerative colitis ^35,36^. Overall, the results of this analysis show that the miRNA profiles of the ileum and colon from matched human NIBD patients are remarkably similar, with only a small set of discriminative miRNAs.

**Figure 2.**
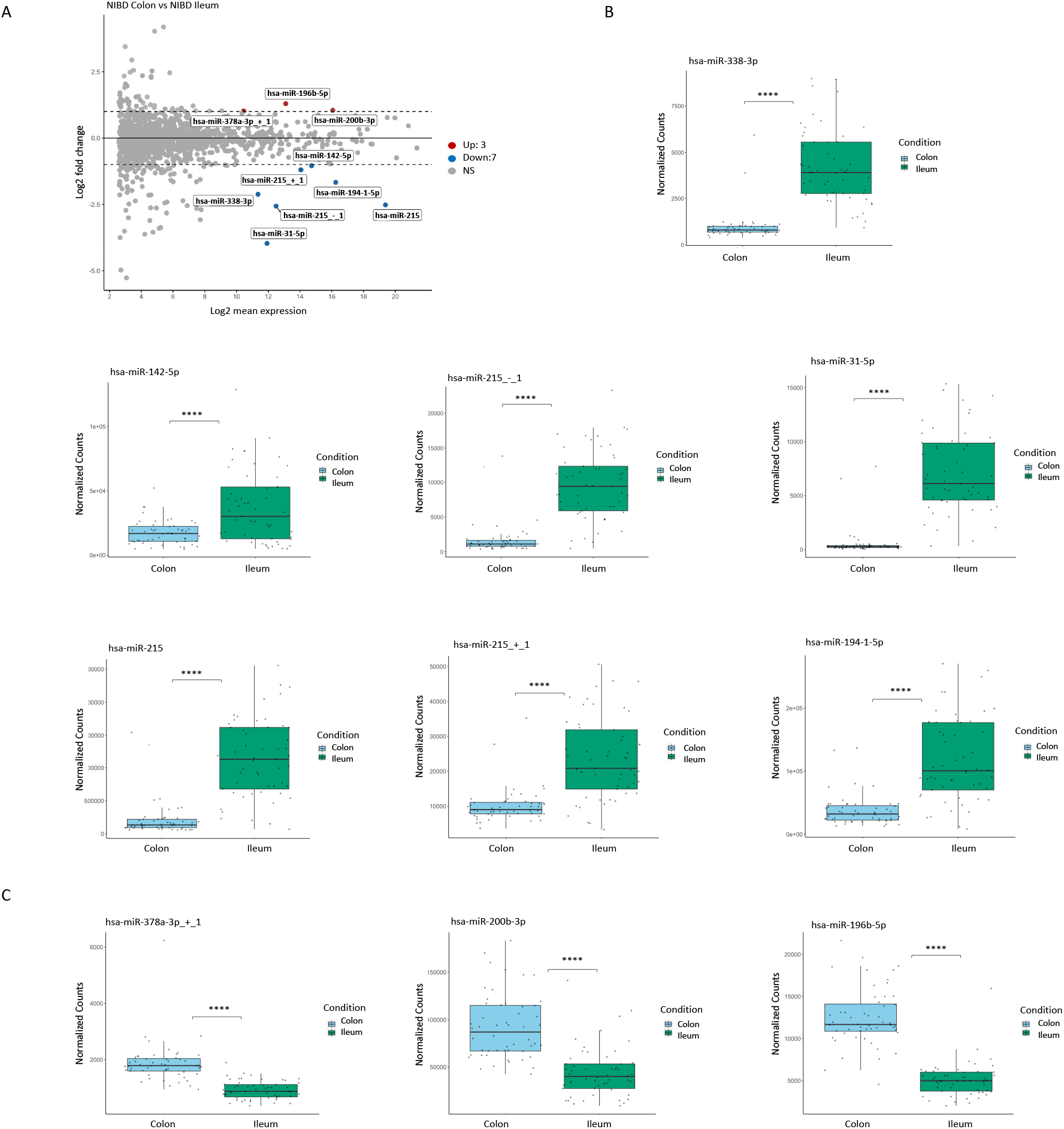
MicroRNA profiles in the colon of pediatric CD separate into two clusters. (A) MA plot showing miRNAs that are significantly differentially expressed between colonic and ileal NIBD patients (n=105) in the colon (left) and ileum (right). Dashed lines represent the log2 fold-change of expression –1.0/+1.0 (horizontal). Up- or downregulated miRNAs are colored red or blue, respectively, with an adjusted p-val < 0.05 and baseMean > 1000. (B) The normalized read counts of seven miRNAs significantly enriched in the ileum and three miRNAs (C) significantly enriched in the colon. Each data point represents a patient sample. (**** p< 0.0001; Student’s t-test).

Next, to identify differentially expressed miRNAs in pediatric CD vs. NIBD, we performed analysis with DESEq2 in both tissue types separately. This analysis revealed 30 significantly altered miRNAs in ileum (12 up, 18 down) and 52 in colon (26 up, 26 down) from pediatric CD patients relative to the corresponding NIBD samples (Figure 3A). Of these, ∼40% are overlapping between the two tissue types (Figure 3B). Although many of the most robustly differentially expressed miRNAs in pediatric CD are shared between ileum and colon, several notable miRNAs are unique to one or the other tissue type. For example, miR-215 and miR-31 are significantly altered in pediatric CD only in ileum and colon, respectively (Figure 3C). Both of these miRNAs are significantly altered in the colon of adult CD patients based on our previously published datasets ^23,25^. Examples of miRNAs found to be significantly altered in both ileum and colon tissue from pediatric CD patients are miR-29b, miR-29c, and miR-375 (Figure 3D).

**Figure 3.**
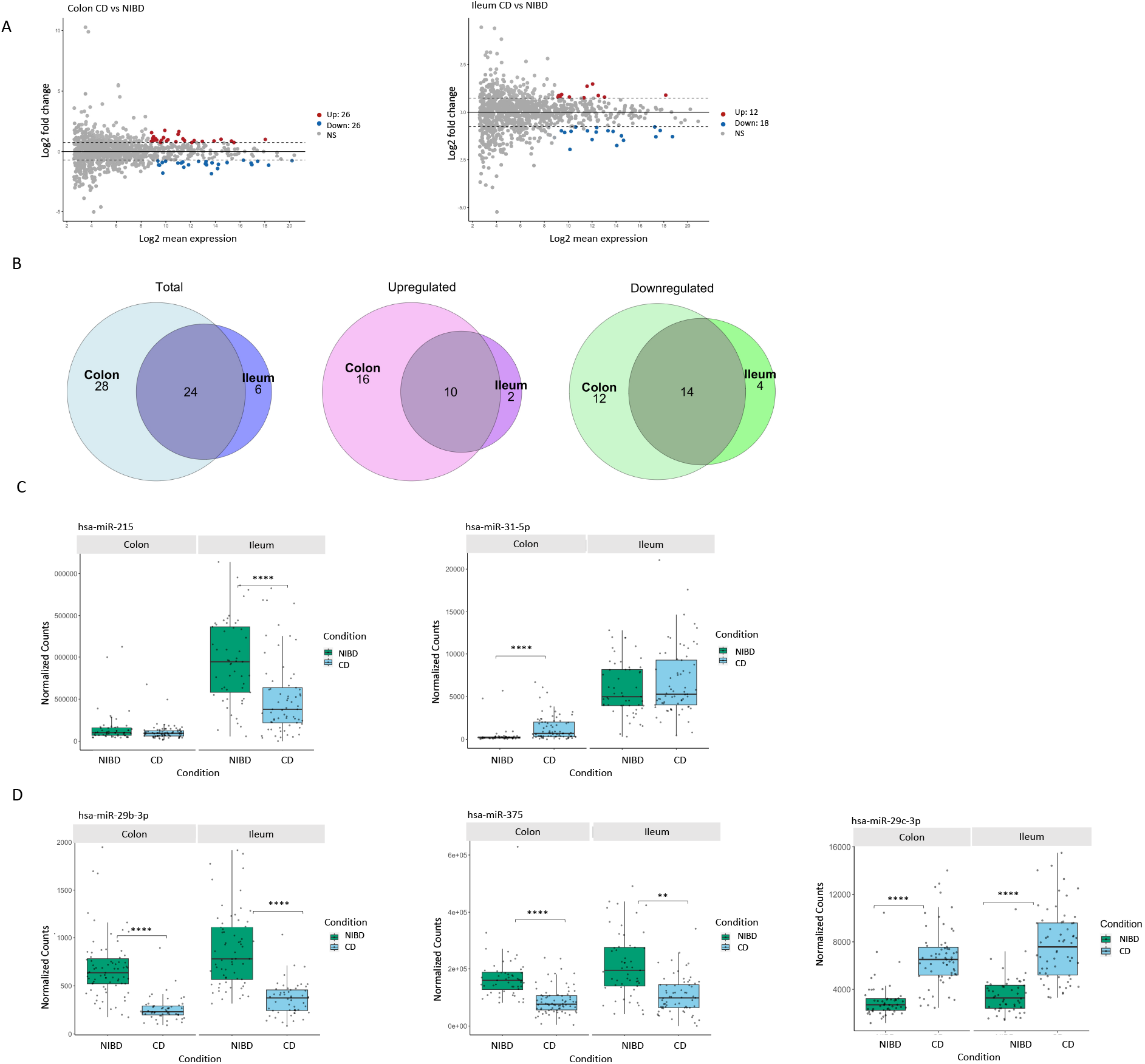
Many microRNAs significantly altered in pediatric CD are shared between the two tissue types. (A) MA plot showing miRNAs that are significantly differentially expressed between CD and NIBD patients (n = 245) in the colon (left) and ileum (right). Dashed lines represent the log2 fold-change of expression –0.75/+0.75 (horizontal). Up- or downregulated miRNAs are colored red or blue, respectively, with an adjusted p-val < 0.05 and baseMean > 500. (B) Venn diagrams of significantly altered pediatric miRNAs (baseMean > 500, p-adj < 0.05, log2FC > 0.75 or <-0.75) in ileum and in colon: total (left), downregulated (middle), and upregulated (right). Paralogs are listed as one miRNA. (C) The normalized counts of miR-215 and miR-31, which are two of the miRNAs significantly differentially expressed specific to the colon or ileal tissue samples. (D) A comparison of normalized counts for miR-29b, miR-375, and miR-29c, which are found to be significantly altered in both ileum and colon tissue from pediatric CD patients. Each data point represents a patient sample. (** p < 0.01, **** p< 0.0001; Wald Test).

### Index levels of ileal miR-29b/c are associated with the development of severe inflammation and stricturing in pediatric CD patients

We sought to determine whether any of the significantly differentially expressed miRNAs in colon or ileum (Figure 3A) are associated with clinical characteristics or predictive of future disease outcomes (Table S1). We first performed binomial regression analysis for all binary outcomes. We found that eight colonic miRNAs are modestly associated with family history; one ileal miRNA isoform (miR-215_-_1) is moderately predictive of surgery with anastomosis; and seven colonic miRNAs (including the miR-21 family and miR-31) are strongly predictive of rectal or sigmoid involvement (Table S2). Both miR-31 and miR-21 have been implicated previously in adult CD as well as in mouse models of colitis. Next, we performed multinomial regression analysis for more complex clinical outcomes and showed that index levels of eight ileal miRNAs (Figure 4A) are significantly associated with the development of at least one of the ileal disease subtypes: severe inflammation, stricturing, or penetrating. Three of these eight are significantly associated with both severe inflammation and stricturing (Figure 4A). Of these three, only two (miR-29b and miR-29c) are not also altered in adult CD based on our previously published analysis (Figure 4B) ^23,25^, suggesting they are prominent and distinguishing features of pediatric CD. We then performed logistic regression analysis to determine whether index levels of miR-29b and miR-29c are associated with the type of ileal disease that a pediatric patient will develop over time (Figure 4C). We found that increasing index levels of ileal (but not colonic) miR-29b and miR-29c are strongly predictive of severe inflammation and stricturing (Figure 4C).

**Figure 4.**
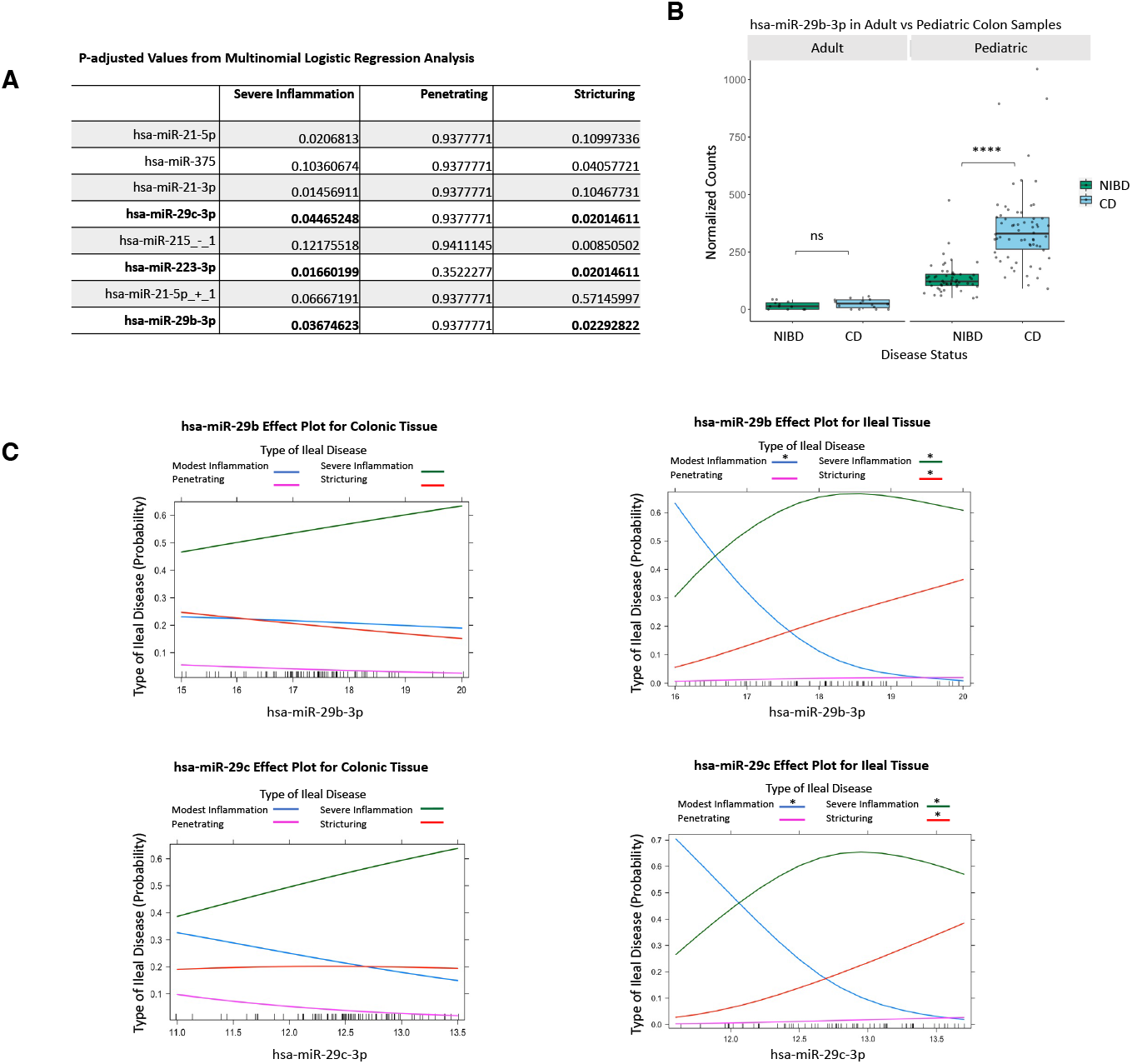
Index levels of ileal miR-29b/c are associated with the development of severe phenotypes in pediatric CD patients. (A) A table of differentially expressed miRNAs in ileal CD vs NIBD with significant p-adjusted values after multinomial logistic regression analysis. Multinomial logistic regression was performed with VST-transformed counts for each miRNA and the type of ileal disease for each patient (modest inflammation used as a reference). FDR was used for multiple testing correction. MicroRNAs in bold have significant p-adjusted values < 0.05 for both stricturing and severe inflammation. (B) Normalized counts of miR-29b in colon tissue from pediatric and CD patients. Each data point represents a patient sample. (C) Effects plots for miR-29b (top) and miR-29c (bottom) with the VST-transformed counts for each microRNA on the x-axis and the probability of association with type of ileal disease on the y-axis. Type of ileal disease is indicated by the colors blue, pink, green, and red for modest inflammation, penetrating, severe inflammation, and stricturing, respectively. Adjusted p-values for each association are placed above the appropriate type of ileal disease. (* p<0.05, **** p< 0.0001; Wald Test).

### Up-regulation of miR-29b is associated with loss of gene encoding tight junction protein PMP22 in mouse and human

To determine the effects of miR-29 up-regulation on the intestine, we leveraged a Dox-inducible miR-29b over-expressing (29OE) mouse model (Methods, Figure 5A). For this study, we believe a whole-body 29OE model is necessary as a starting point because the specific cell types or even tissue layers in the intestine in which miR-29 up-regulation occurs in pediatric CD is not known. We first performed histological analysis of duodenal, jejunal, and ileal tissue isolated after 60 days of post-natal Dox administration (29OE/+Dox) and compared to control mice (29OE/-Dox). In two separate rounds of analysis (each with n=4 29OE/+Dox and n=4 control mice), we did not observe any gross disturbances in small intestinal architecture nor any substantial differences in intestinal crypt depth or density in 29OE/+Dox relative to control (Figure S3).

**Figure 5.**
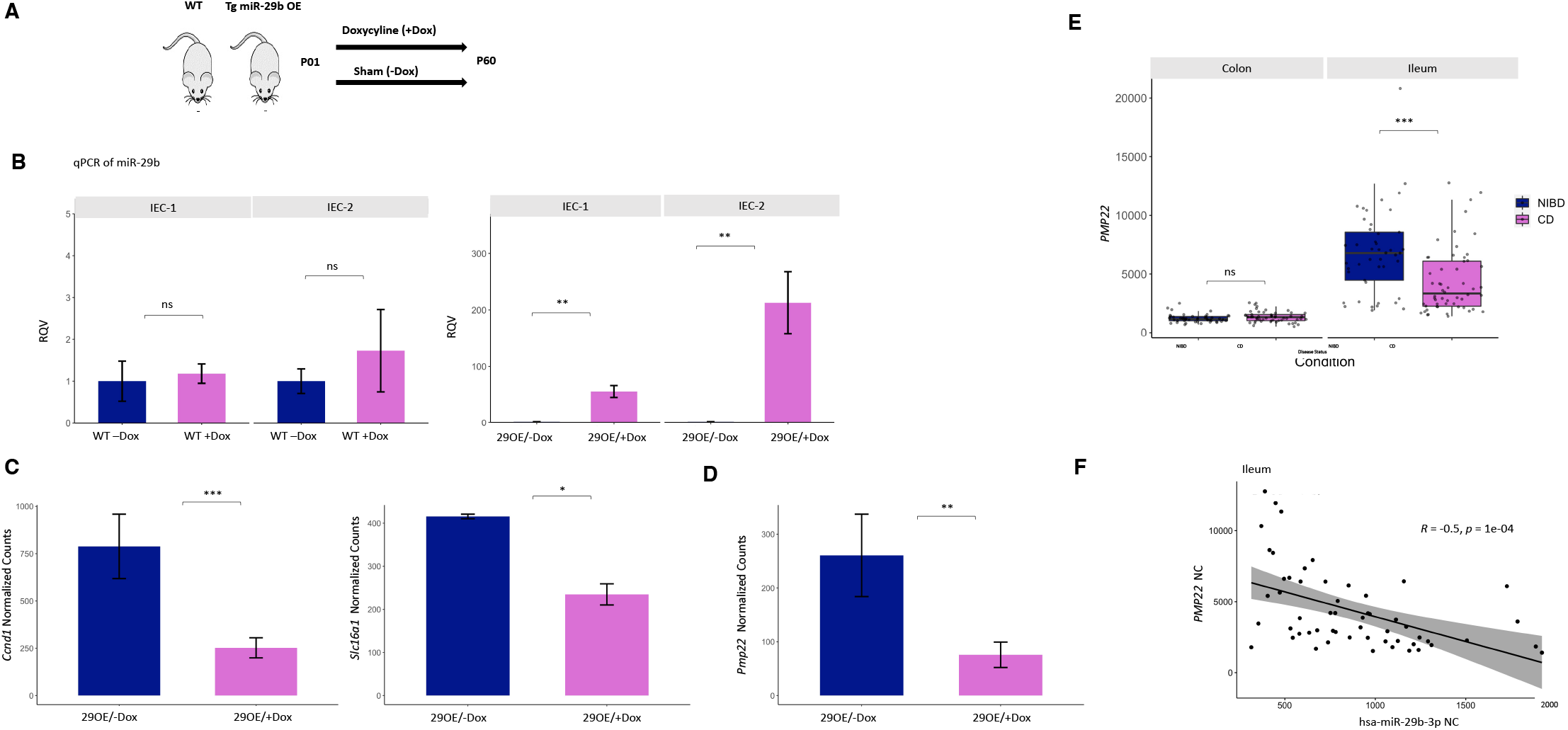
Upregulation of miR-29b is associated with loss of PMP22 in both mouse and human. (A) Schematic of doxycycline-inducible miR-29b over-expressing (29OE) mouse model. (B) qPCR showing the relative quantitative value (RQV) of miR-29b in 29OE mice without doxycycline treatment (29OE/-Dox, n=4) (left) and with doxycycline treatment (29OE/+Dox, n=4) for IEC-1 and IEC-2 (right). IEC-1 and IEC-2 represent two independent intestinal epithelial fractions from the same mice. Dark blue represents the –doxycycline treatment and pink represents the +doxycycline treatment. (C) Normalized counts of previously validated miR-29 target genes *Ccnd1* and *Slc16a1* in the transgenic mouse model. Dark blue represents the –doxycycline treatment (n=3) and pink represents the +doxycycline treatment (n=3). (D) Normalized counts of miR-29 predicted target gene, *Pmp22*, in the transgenic mouse model. Dark blue represents the –doxycycline treatment (n=3) and pink represents the +doxycycline treatment (n=3). (E) Normalized counts of miR-29 predicted target gene *PMP22* in pediatric CD and NIBD in both tissue types. (F) Correlation of miR-29b (x-axis) and *PMP22* (y-axis) in pediatric ileum samples. Data presented as mean ± SEM (* p<0.05, ** p<0.01, *** p<0.001; Student’s t-test).

Since a compromised epithelial barrier is one of the hallmark features of CD, we next isolated intestinal epithelial cells (IECs) and confirmed that miR-29b levels are significantly elevated in 29OE/+Dox mice relative to control (Figure 5B), and that this induction is dependent upon the miR-29 over-expression cassette and not due to Dox treatment alone (Figure 5B).

RNA-seq analysis of IECs in 29OE/+Dox relative to control showed significant down-regulation of 72 genes, including very well-established miR-29 target genes such as *Ccnd1* and *Slc16a1* (Figure 5C)^37,38^. These data confirm at the gene expression level the expected gain-of-function of miR-29 in the intestinal epithelium in the 29OE/+Dox mice.

Notably, we also found that *Pmp22*, a predicted miR-29 target gene, is among the most significantly down-regulated genes in 29OE/+Dox mice (Figure 5D). This gene encodes a tight junction protein that very recently was shown to promote intestinal barrier function^39^. Although miR-29 has been implicated in the control of barrier capacity through regulation of the tight junction protein Cldn1 previously, *Pmp22* has not been reported as a miR-29 target in the intestine ^40^.

To determine whether this regulatory relationship holds in human, we analyzed our previously reported RNA-seq data from a majority subset of the same human samples used in this study (n=203) (Figure S4). We found that *PMP22* is among only 16 genes that are significantly down-regulated in both the ileum of pediatric CD patients (relative to NIBD controls) and in IECs of 29OE/+Dox mice (relative to 29OE/-Dox controls) (Table S3). Moreover, we observed that *PMP22* is much more highly expressed in pediatric human ileum compared to colon (Figure 5E) and also significantly suppressed only in the ileum and not the colon of pediatric CD compared to NIBD patients (Figure 5E). Notably, we also observed that miR-29b levels are highly significantly inversely correlated with *PMP22* in the ileum of pediatric CD patients (Figure 5F). Taken together, these data point to a new miR-29 target in the small intestine, the reduction of which in the context of miR-29 up-regulation may contribute to the compromised barrier observed in pediatric CD.

### Over-expression of miR-29b in mice leads to dramatic reduction of Paneth cell gene markers

Upon further analysis of the murine RNA-seq data from the jejunum, we observed dramatic down-regulation of six major Paneth cell markers (Figure 6A). Contrastingly, we observed only modest effects on goblet and enteroendocrine cell markers (Figure 6A), and very little influence on stem or enterocyte markers (Figure 6A). We then measured by real time quantitative PCR (RT-qPCR) the levels of marker genes of intestinal stem cells (*Lgr5*) and four different major lineages of the intestinal epithelium (enterocyte, goblet, enteroendocrine, and Paneth). The most dramatic effect was observed for a classic marker of crypt-based Paneth cells, lysozyme 1 (*Lyz1*), downregulated by more than 20-fold in 29OE/+Dox relative to control (Figure 6B). No significant change in *Lyz1* or any other marker was detected in wild-type mice (without the miR-29b over-expression cassette) treated with Dox for the same duration of time (Figure 6B). RT-qPCR for additional Paneth cell markers, *Defa17*, miR-152, and *Copz2*, in IECs revealed a similar down-regulation in 29OE/+Dox relative to control (Figure 6C-E). The latter two are particularly informative as they are not thought to be physically associated with granules, unlike Defa17 and Lyz1, suggesting that there is a loss of Paneth cells and not merely a granulation defect.

**Figure 6.**
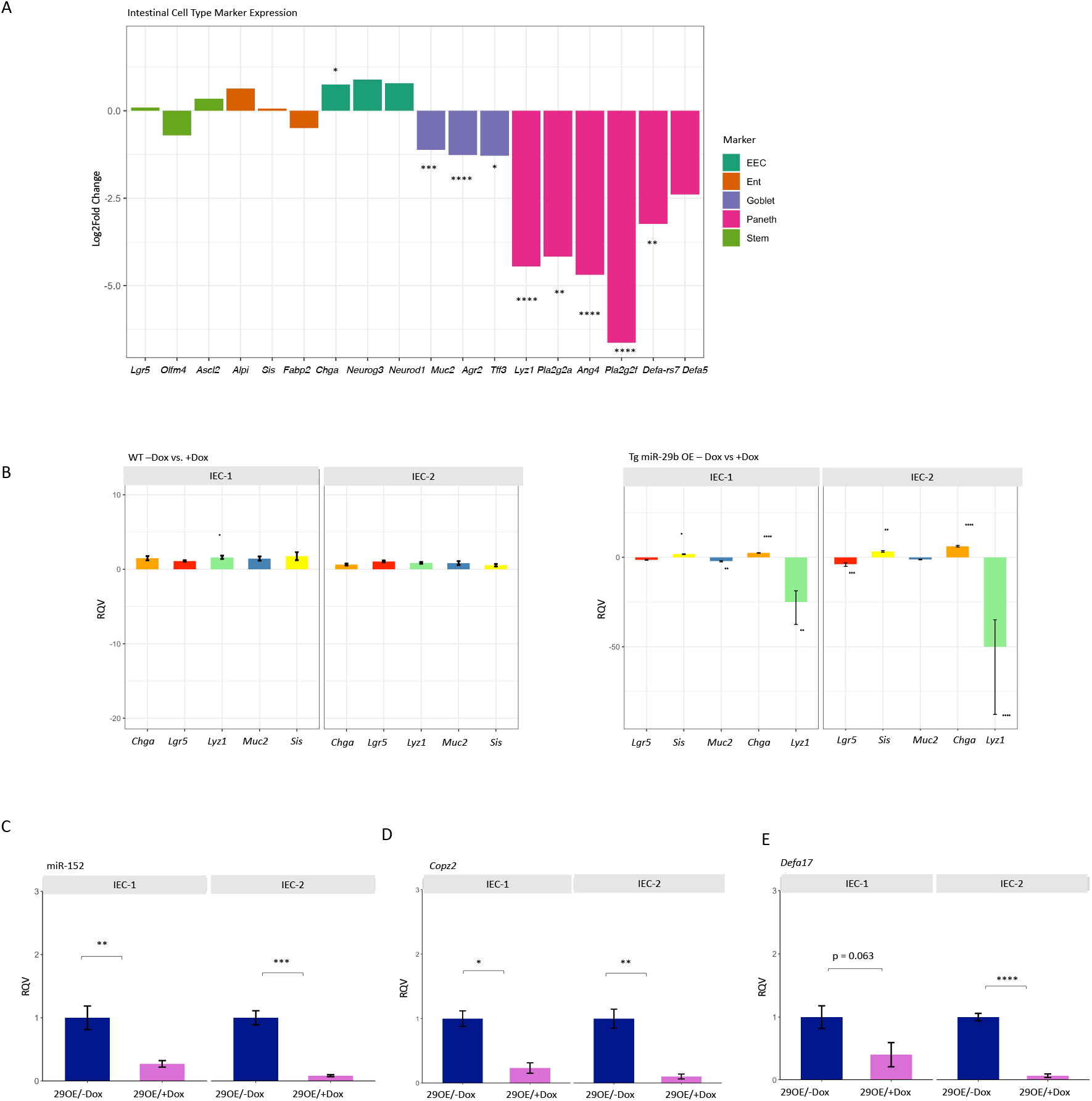
Over-expression of miR-29b in mice leads to dramatic reduction of Paneth cell gene markers. (A) RNA-seq data from the jejunum shows log2 fold-change (y-axis) of intestinal epithelial cell type markers (x-axis) in 29OE/+Dox (n=3) vs. 29OE/-Dox (n=3). The markers for each cell type are colored in green, orange, teal, purple, and pink for the following cell types: stem cell, enterocyte, enteroendocrine cell, goblet cell, and Paneth cell, respectively. ((* p <0.05, ** p< 0.01, *** p < 0.001, **** p < 0.0001; Wald Test) (B) qPCR of 29OE/-Dox (left, n=4) and 29OE/+Dox (right, n=4) showing the relative quantitative value (RQV) for marker genes of intestinal stem cells (Lgr5) and four different major lineages of the intestinal epithelium (enterocyte, goblet cell, enteroendocrine cell, and Paneth cell) with and without doxycycline treatment for IEC-1 and IEC-2. qPCR showing the relative quantitative value (RQV) of miR-152 (C), *Copz2* (D), and *Defa17* (E) 29OE for IEC-1 (n=4,4) and IEC-2 (n=4,4). Dark blue represents the –doxycycline treatment and pink represents the +doxycycline treatment. Data presented as mean ± SEM (* p <0.05, ** p< 0.01, *** p < 0.001, **** p < 0.0001; Student’s t-test).

### Gain of miR-29b leads to loss of Paneth cells in mice

Matching the results from the transcriptomic study, H&E analysis showed that the number of granulated cells (which we use as a proxy for Paneth cells) per crypt is significantly reduced in 29OE/+Dox mice compared to 29OE/-Dox controls (Figure 7A). We next performed Lyz1 immunofluorescence (IF) analysis, which showed an even more dramatic loss of canonical Paneth cells (Figure 7B). H&E and IF analyses in an independent cohort of mice confirmed these results (Figure S5). Alcian blue staining revealed only a comparatively modest effect of miR-29b over-expression on goblet cell number in both crypts (Figure 7C) and villi (Figure 7D), consistent with the results of the gene marker analysis (Figure 6A). These findings were not observed in the H&E analysis for wild-type mice (without the miR-29b over-expression cassette) treated with Dox for the same duration of time (Figure S6).

**Figure 7.**
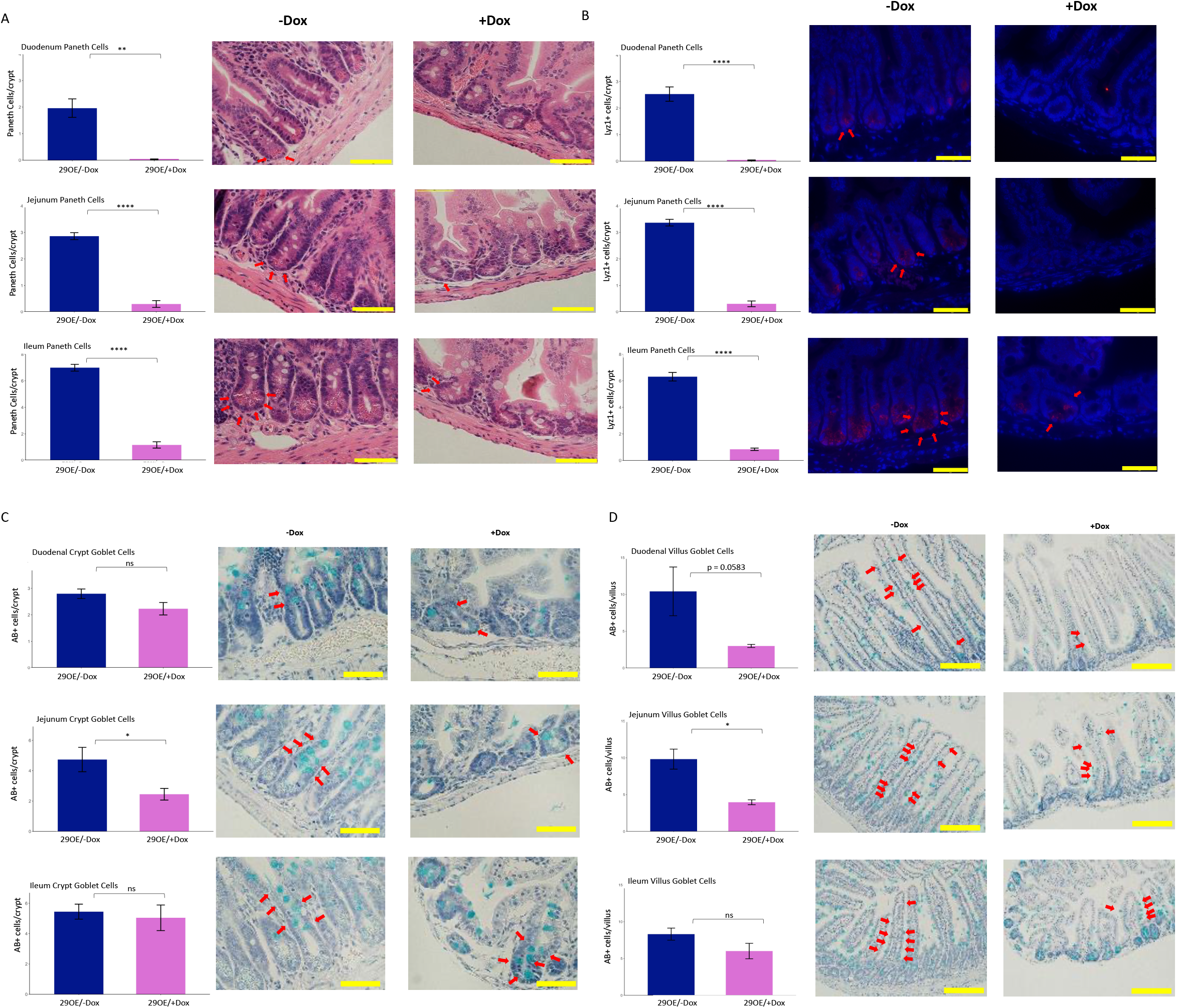
Gain of miR-29b leads to loss of Paneth cells. (A) Paneth cell counts per crypt from brightfield H&E-stained tissue sections of 29OE/-Dox (n=4) and 29OE/+Dox (n=4) in the proximal duodenum, mid-jejunum, and distal ileum (x600 images). (B) Paneth cell counts per crypt from Lyz1 immunofluorescent (red) and DAPI fluorescent (blue) tissue sections of 29OE/– Dox (n=4) and 29OE/+Dox (n=4) in the proximal duodenum, mid-jejunum, and distal ileum (x600 images). (C,D) Goblet cell counts per crypt (C) or per villus (D) from Alcian blue and eosin–stained tissue sections of 29OE/–Dox (n=4) and 29OE/+Dox in the proximal duodenum, mid-jejunum, and distal ileum (x600 images for crypts, x200 images for villi). Yellow scale bars represent 50 µm at x600 and 100 μm at x200. Individual Paneth and goblet cells are indicated by red arrows. Dark blue represents the –doxycycline treatment and pink represents the +doxycycline treatment. Data presented as mean ± SEM (* p <0.05, ** p< 0.01, **** p < 0.0001; Student’s t-test).

### miR-29b/c levels are linked to Paneth cell number in pediatric CD patients

Based on the functional studies in mice, we hypothesized that miR-29b/c levels are correlated with Paneth cell number in pediatric CD patients. To test this hypothesis, we first selected the patients with the highest or lowest levels of miR-29b/c, termed High-29 (n=20) or Low-29 (n=19), respectively (Figure 8A). Among these, 9 samples dropped out of further analysis due to not meeting our histology criterion for displaying at least 10 well-oriented crypts with fully discernible crypt bottoms. Of the remaining 30 samples, 19 (High-29, n=10; Low-29, n=9) were analyzed for inflammation. We found that miR-29b levels are significantly correlated with inflammation score (Figure 8B). High-29 samples were found to have significantly reduced levels of *DEFA5* and *DEFA6*, which are specific markers of Paneth cells (Figure 8C, Table 3). The samples (n=30) were then subject to H&E analysis, which revealed that the High-29 group is associated with significantly fewer Paneth cells per crypt (Figure 8D-E). Taken together with the previous results, these findings are strongly indicative of a dominant regulatory effect of miR-29b on Paneth cells in mice and humans.

**Table 3.** Demographics of pediatric CD patients from RNA-sequencing analysis for High-29b and Low-29b categories.

| Patient Characteristics | Low (Average) | High (Average) |
| --- | --- | --- |
| Diagnosis Age (in years) | 9.33 | 13 |
| Grouped Ages |  |  |
| VEO (<6) | 2 | 0 |
| Child (7-12) | 9 | 4 |
| Teen (13-17) | 4 | 9 |
| Sex |  |  |
| Female | 6 | 7 |
| Male | 9 | 6 |

**Figure 8.**
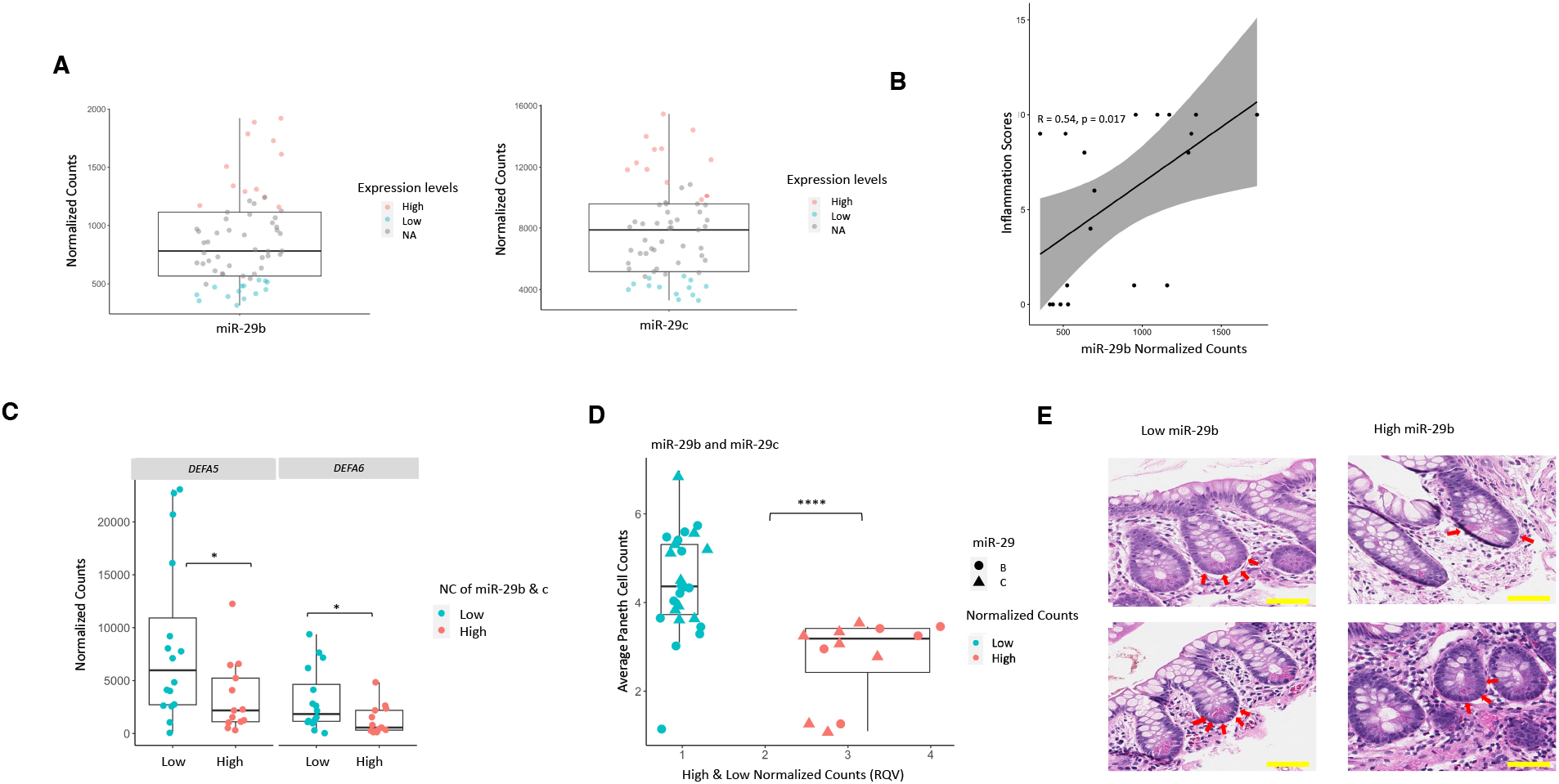
High levels of miR-29b/c are linked to low numbers of Paneth cells in pediatric CD patients. (A) The normalized counts of miR-29b/c in the selected High-29 (n=20) or Low-29 (n=19) groups. Each data point represents a patient sample. Red dots represent the patients in the High-29 group and blue dots represent those in the Low-29 group, the remaining patients are presented in gray. (B) Correlation of miR-29b (y-axis) and inflammation scores (x-axis) across patients in the pediatric High-29 and Low-29 groups (High-29, n=10; Low-29, n=9). (C) Normalized counts of miR-29b/c in High-29 and Low-29 samples for both *DEFA5* and *DEFA6*. (D) Relative quantitative values (RQV) of miR-29b/c expression in High-29 and Low-29 samples (x-axis) and average Paneth cell counts per crypt (y-axis). (E) Two representative brightfield H&E-stained images of ileal crypts from High-29 and Low-29 patients (x600). Yellow scale bar represents 50 μm. Each data point represents a patient sample. (* p< 0.05; **** p< 0.0001; Student’s t-test).

## Discussion

To our knowledge, this study represents the first comprehensive miRNA analysis of a large cohort of pediatric CD patients. Major strengths and novelties of this study include: (1) large sample size of treatment-naïve patients (n=245); (2) patient-matched ileal and colonic tissue; (3) detailed regression analysis with clinical characteristics; (4) discovery of index biopsy miRNA indicators of disease outcomes; and (5) mouse and human studies linking aberrant miR-29 with the loss of Paneth cells and altered *PMP22* expression. A primary finding is that miR-29 and possibly other miRNAs could be used as prognostic indicators of disease subtype and/or severity. Major open questions that our study does not address include: (1) how is miR-29 mis-regulated in pediatric CD?; (2) are the effects of miR-29 over-expression on Paneth cells a developmental phenotype?; and (3) in what intestinal cell type(s) or mucosal layer(s) is miR-29 over-expression the most dominant and functionally relevant in the context of pediatric CD?

### These questions warrant further investigation in follow-up studies

The answers to the third question listed above will be particularly valuable for determining the downstream targets of miR-29 that are most critical in the context of pediatric CD. In the murine RNA-seq data presented in this study, none of the genes encoding canonical regulators of Paneth cell differentiation^41^ (*Sox9, Atoh1, Erbb3*), appear to be affected in the intestinal epithelium upon miR-29 over-expression. Intriguingly, though, we do observe a dramatic decrease in *Ciita*, which codes for the master transcriptional regulator of MHC Class II genes ^42^. *Ciita* is a predicted target of miR-29 and loss of MHC Class II activity specifically in intestinal stem cells can lead to significantly reduced secretory cell allocation ^43^. Taken together, these data raise the possibility that up-regulation of miR-29 leads to increased direct targeting and suppression of *Ciita*, which in turn reduces MHC Class II signaling and Paneth cell differentiation. This hypothesis requires extensive rigorous evaluation that is well outside the scope of the present study, but we believe it merits future investigation. It is also quite possible that the up-regulation of miR-29 in pediatric CD is greatest in cells from layers beneath the epithelium, including fibroblasts, lymphatic endothelial cells, telocytes, immune cells, and/or enteric neurons. For example, one possibility is that increased miR-29 specifically in lamina propria T cells leads to reduced *DNMT3A*, a well-established miR-29 direct target, which would decrease promoter methylation of the interferon-gamma gene (*IFNG*), increase IFNg levels, and promote chronic inflammation^44,45^. We propose that future studies should focus on determining the cell types driving the aberrant miR-29 signal in pediatric CD in order to uncover direct targets and molecular mechanisms that underpin the association of miR-29 with inflammation, stricturing, and/or Paneth cell loss.

miR-29 has received attention previously as a potent regulator of several gastrointestinal phenotypes^15^. A few different studies of irritable bowel syndrome (IBS) have shown that miR-29 promotes gut permeability in mice by directly targeting and suppressing tight junction proteins ^40,46^. In this context, up-regulation of miR-29 would severely compromise the gut barrier, and thereby promote IBS disease severity. In other work, though, it has been demonstrated that miR-29 suppresses intestinal fibrosis and therefore could be a protective factor in IBD ^47,48^. Moreover, another study has shown that loss of miR-29 may exacerbate inflammatory phenotypes in the intestine ^49^. In fact, at least one study has suggested that a miR-29 mimic is a potential therapeutic for IBD, especially in cases of severe inflammation ^50^. These separate reports of anti-inflammatory, anti-fibrotic, and yet barrier-compromising functions paints a highly multifaceted picture of miR-29 in the gut. As it pertains to IBD, it is possible that miR-29 has both antagonistic and protective functions depending on the cellular context, disease etiology, age of onset, and/or time point during disease progression.

Our discovery that miR-29 may also suppress Paneth cells provides another means by which gut permeability and inflammation may increase. Specifically, we suggest that an early increase in miR-29 leads to Paneth cell loss, which dampens anti-microbial activity, likely promotes small intestinal dysbiosis, which in turn compromises barrier integrity leading to increased risk of inflammation. Paneth cell defects have long been implicated in the pathogenesis of CD, and in fact are more prevalent in children compared to adults ^51,52^ We suggest that the aberrant elevation of miR-29 in pediatric CD, but not adult CD, may contribute to this difference. Our findings add to the rich and complex web of intestinal miR-29 behavior.

Several reports have called for the investigation of both miR-29 mimics and inhibitors as potential therapeutics in IBD and related chronic disorders of the gut. However, we strongly urge caution, as miR-29-based therapeutics are likely to be challenging given the context-specific functions of miR-29 described above. Further work is needed to sort out the regulatory effects of miR-29 in distinct cell types of the intestine during different stages of disease progression. We are even more intrigued by the potential of miR-29-based therapeutics for acute gut conditions (such as microbial infections), or conditions such as necrotizing enterocolitis, for which Paneth cell defects are critical to pathogenesis. For example, is miR-29 strongly up-regulated after *Salmonella* or *Listeria* infection, and if so, could that be responsible at least in part for the Paneth cell defects reported under those conditions? We believe that such questions, while outside scope of the present study, merit detailed investigation in the future.

The mechanistic underpinnings of pediatric CD are still so poorly understood. We believe this study marks an important turning point in the investigation of miRNAs in pediatric CD and provides a rich resource to the research community for the identification of key regulators of the disease, well beyond miR-29. For example, our data points to a significant loss of miR-375 in CD. At least one prior study showed that the loss of miR-375 up-regulates pro-inflammatory factors such as TLR4 and NFKB ^53^. We believe that more detailed investigation of the mechanisms by which miR-375, and other miRNAs revealed by this dataset, might control the inflammatory, stricturing, or other IBD phenotypes is warranted. Such studies may uncover novel and effective therapeutic avenues for pediatric and/or adult CD.

## Materials and Methods

### Patient Population and sample acquisition

Patient samples were acquired from the University of North Carolina Multidisciplinary IBD Center, abiding by institutional review board-approved protocols (Study ID#: 15-0024). All samples were procured from treatment-naive pediatric patients undergoing endoscopy for suspicion of IBD between 2008 and 2012. Non-IBD control samples were defined as patients with biopsies that were subsequently identified as histologically normal. Clinical information gathered in this study include demographic and clinical variables: family history, age, sex, disease duration, disease location. Information from patient clinical parameters is provided (Table S1). From FFPE tissue, mucosal biopsies were obtained during colonoscopy from macroscopically unaffected sections of the ascending colon and terminal ileum at the time of surgery. These index biopsies were also confirmed by a pathologist (D.G. Trembath) to have no active inflammation.

### Small RNA library preparation and sequencing

RNA was isolated from FFPE tissue using the Roche High Pure miRNA Isolation Kit following the manufacturer’s protocol as previously described ^25^. RNA purity was quantified with the NanoDrop 2000 instrument (Thermo Fisher Scientific, Waltham, MA), and RNA integrity was quantified with the Agilent 2100 Bioanalyzer (Aglient Technologies, Santa Clara, CA). Small RNA libraries were generated using the TruSeq Small RNA Sample Preparation Kit (Illumina, San Diego, CA). Sequencing (Single-end, 50bp) was performed on the HiSeq 2500 platform (Illumina, San Diego, CA) at the Genome Sequencing Facility of the Greehey Children’s Cancer Research Institute (University of Texas Health Science Center, San Antonio, TX). Previously published raw sequencing data can be accessed through GEO accession no. <u>GSE101819</u>. Additional samples will be uploaded through GEO at a future date.

### Small RNA-sequencing analysis

Read quality was assessed using FastQC. miRquant 2.0^54^ was used for read trimming, miRNA annotation, and quantification. Briefly, reads were trimmed using Cutadapt, then aligned to the human genome (hg19) using both two different mapping tools (Bowtie^55^ and SHRiMP^56^), and raw miRNA counts were quantified and normalized using DESeq2^57^ to determine significance. Samples with fewer than 1 million reads mapping to miRNAs were removed from further analysis.

Both grouped age (VEO < 6, Child = 7-12, Teen = 13-17), SIM, and sex of the pediatric samples were accounted for using the limma^58^ function removeBatchEffect. After correcting for covariates, PCA analysis was performed using the log normalized DESeq2^57^ values and the plotPCA^57^ function in R. Hierarchical clustering was performed using the log normalized DESeq2 values. Using the stats function dist, the Euclidean distance between samples was computed based on expression and plotted with pheatmap. Individual PCAs were computed for both colonic and ileal tissue types using log normalized DESeq2^57^ values with SIM, grouped age, and sex accounted for as covariate. MA plots were created after correcting for covariates and using ggmaplot from ggpubr^59^ with parameters set for baseMean, log2FC, and padj.

### Regression analysis

Associations of differentially expressed miRNAs from both colonic and ileal tissue with binary clinical parameters were explored using generalized linear models (GLM). With categorical variables from clinical data, probabilities were obtained using the multinom function from “nnet”^60^ that produced fitted values from a multinomial regression model. Multiple testing correction was performed using FDR adjustment.

### Mouse models

The miR-29 over-expression mouse model used in this study was first described in the following publication: https://www.biorxiv.org/content/10.1101/2022.11.29.518429v1.

### Jejunal epithelial cell preparation

Harvested small intestine from doxycycline exposed and unexposed WT and miR-29b overexpressing mice was measured and divided into three equal segments. The middle region was considered jejunum. Subsequent to luminal flushing with ice cold phosphate buffered saline (PBS), the tissue was longitudinally cut and subjected to incubation in 3 mM EDTA in ice cold PBS with 1% (v/v) primocin (InvivoGen, San Diego, CA) for 15 min at 4°C. The mucosa of the intestinal pieces was gently scrapped of mucus, shaken in ice cold PBS with 1% (v/v) primocin (InvivoGen) for 2 minutes, and incubated in fresh 3 mM EDTA in ice cold PBS with 1% (v/v) primocin (InvivoGen) for 40 min at 4°C. After 2 to 6 minutes of gentle manual shaking in ice cold PBS with 1% (v/v) primocin (InvivoGen), the intestinal pieces were inspected microscopically (x100 magnification) for detached intestinal crypts and villi, and then diluted 1:2 with ice cold PBS with 1% (v/v) primocin (InvivoGen). Material that filtered through a 70 μm cell strainer was collected and referred to as jejunal epithelial cell fraction one (IEC-1), while material that was collected with washing of the cell strainer surface with ice cold PBS with 1% (v/v) primocin (InvivoGen) was referred to as jejunal epithelial cell fraction two (IEC-2). IEC-1 and IEC-2 preparations were then pelleted by centrifugation at 110 x g for 10 min at 4°C. For RNA extraction, collected pellets were resuspended in 200 μL of lysis buffer (Buffer RL, Norgen Biotek, Thorold, ON, Canada), vortexed for 10 s, and stored at −80°C.

### RNA extraction and real-time qPCR

Total RNA was isolated using the Total RNA Purification kit (Norgen Biotek). TaqMan microRNA Reverse Transcription kit (Life Technologies) was used for reverse transcription of miRNA. High Capacity RNA to cDNA kit (Life Technologies, Grand Island, NY) was used for reverse transcription of RNA for gene expression analysis. Both miRNA and gene expression qPCR were performed using TaqMan assays (Life Technologies) with either TaqMan Universal PCR Master Mix (miRNA qPCR) or TaqMan Gene Expression Master Mix (mRNA qPCR) per the manufacturer’s protocol on a BioRad CFX96 Touch Real Time PCR Detection System (Bio-Rad Laboratories, Richmond, CA). Reactions were performed in duplicate or triplicate using either U6 (miRNA qPCR) (Life Technologies, Assay ID: 001973) or Rps9 (mouse mRNA qPCR) (Life Technologies, Assay ID: Mm00850060_s1) as the normalizer. miRNA expression was assayed with use of the following probes: hsa-miR-29b (Life Technologies, Assay ID: 000413) and hsa-miR-152 (Life Technologies, Assay ID: 000475). Gene expression was assayed with use of the following probes: *Chga* (Life Technologies, Assay ID: Mm00514341_m1), *Copz2* (Life Technologies, Assay ID: Mm04203911_m1), *Defa17* (Life Technologies, Assay ID: Mm04205962_gH), *Lgr5* (Life Technologies, Assay ID: Mm00438890_m1), *Lyz1* (Life Technologies, Assay ID: Mm00657323_m1), *Muc2* (Life Technologies, Assay ID: Mm01276696_m1), and *Sis* (Life Technologies, Assay ID: Mm01210305_m1). Any undetectable samples were not included in the analysis.

### Mouse RNA library preparation, sequencing, and analysis

RNA-sequencing libraries were prepared using the total extracted RNA from the IEC-1 preparations of doxycycline exposed and unexposed miR-29b overexpression mice. RNA was quantified with the NanoDrop 2000 (Thermo Fisher Scientific, Waltham, MA) and RNA integrity was assessed by the Agilent 4200 Tapestation (Agilent Technologies, Santa Clara, CA). Libraries were prepared by the Cornell TREx facility using NEBNext Ultra II Directional Library Prep Kit (New England Biolabs, Ipswich, MA) with ribosomal RNA depletion. Sequencing was performed on the NextSeq500 platform (Illumina, San Diego, CA) at the Genomics Facility in the Biotechnology Research Center at Cornell University. Raw sequencing data is available through GEO accession [GEO Accession Number will be added at a future date]. Read quality was assessed using FastQC. RNA-seq data were mapped to the mm6 genome with STAR^61^.Transcripts were quantified with Salmon^62^ using GENCODE release 25 transcript annotations. Normalization and differential analysis was then performed using DESeq2^57^.

### Human RNA library preparation, sequencing, and analysis

Total RNA was isolated from FFPE tissue using Quick-RNA FFPE MiniPrep (Zymo Research, Irvine, CA). Purification was performed using the MagMAX kit in the KingFisher system (Ther-moFisher, Carlsbad, CA). Sequencing libraries were then prepared through the TruSeq Stranded Total RNA with Ribo-Zero (Illumina, San Diego, CA). The NovaSeq 6000 platform was used for paired-end (50 bp) sequencing (Illumina, San Diego, CA). Salmon^62^ was then used to quantify transcripts. Samples with low transcript numbers (<25,000) and poor transcript integrity numbers (TIN) were eliminated from further analysis (n =2). Samples that failed to cluster with their respective tissue type (ileum or colon) through principal component analysis (PCA) were also discarded from analysis (n = 5).

PCA analysis accounted for the three covariates that contributed to the greatest variation among samples (batch, sex, and TIN). RUVSeq^63^ identified additional unwanted variation by accounting for the top 1000 genes with the lowest variance out of the top 5000 genes with highest variance, these were dictated as the control genes. Through this analysis it was determined that one factor of unwanted variation should be used in the final analysis, due to the observed variation in DEGs identified by DESeq2^57^, relative log expression plots, and correlation between factors of unwanted variation and the outcome.

### Mouse tissue histology and histological analysis

Mouse proximal duodenal, mid-jejunal, and distal ileal tissue were fixed in 4% (v/v) neutral-buffered paraformaldehyde, embedded in paraffin, and cut into 5 μm transverse sections for various staining experiments. Haemotoxylin and eosin (H&E) staining was performed for morphometric analyses (crypt depth, villus height, and crypt density) and Paneth cell count determination. Alcian blue (pH 2.5) and eosin staining (AB&E) was performed for goblet cell count determination. Paneth cell counts were also determined by immunofluorescent staining of lysozyme. Briefly, after deparaffinization and antigen retrieval with citrate buffer (10 mM citric acid, 0.05% (v/v) Tween 20, pH 6.0), sections were blocked with 10% (v/v) normal goat serum in PBS for 1 h at room temperature, incubated with rabbit anti-LYZ primary antibody (1:1000, Abcam, Waltham, MA, cat. ab108508) in PBS with 0.1% (w/v) bovine serum albumin overnight at 4° C, followed by goat anti-rabbit Alexa fluor 594 secondary antibody (1:1000, Invitrogen, Carlsbad, CA, cat. A1102) incubation in PBS with 0.1% (w/v) bovine serum albumin for 1 h at room temperature. 4’,6-diamidino-2-phenylindole (DAPI) (0.1 mg/mL in PBS, Invitrogen, cat. D1306) was used to visualize nuclei. Images were captured using a BX53 Olympus scope (Olympus, Center Valley, PA). Paraffin embedding, sectioning, and tissue staining with H&E and AB&E were performed by the Animal Health Diagnostic Care Histology Laboratory at Cornell University. Images were analyzed for histomorphometric measurements and cell counts with ImageJ software. At least 10 intact, well-sectioned crypts and 10 intact, well-sectioned villi were used for acquiring histomorphometric measurements and cell counts.

### Human patient tissue histology and histological analyses

Two pathologists (LCZ and SB) independently and blindly graded the inflammatory activity (x400) of ileal H&E stained sections using the following criteria. High degree of inflammation: neutrophilic activity on 7-10/10 high power fields; Intermediate degree of inflammation: neutrophilic activity on 3-6/10 high power fields; low degree of inflammation: neutrophilic activity on 0-2/10 high power fields. Using brightfield images at x600 magnification, the crypt base eosinophilic granulated Paneth cells of at least 10 well-oriented crypts with fully discernible crypt bottoms were counted.

### Statistics

R software version 4.1.0 was used for these data analyses. All small RNA annotation and quantification was conducted through miRquant^54^. RNA-seq and small RNA-seq data were analyzed for differential expression using DESeq2^57^, with p-values FDR adjusted for statistical significance. Significance of differential expression in RT-qPCR experiments was assessed using Student’s t test (unpaired, 2-tailed) to compare 2 groups of independent samples. Multinomial logistic regression analysis also was performed using R software and p-values were adjusted using FDR. All statistical tests used are detailed in the figure legends. P-values < 0.05 are considered statistically significant. NS = not significant, * = p<0.05, ** = p<0.01, *** = p<0.001. In figure panels, unless otherwise noted, quantitative data are reported as an average of biological replicates ± standard error of the mean for all mice studies. For human samples, quantitative data are reported with standard deviation.

## Figure Legends

**Supplemental Figure 1**. Unsupervised hierarchical clustering of the Euclidean distances among patient-matched (n=228) pediatric samples was calculated based on VST normalized counts. The analysis for the colon (left) and ileal tissue (right) account for the covariates of small-RNA integrity metric (SIM), grouped ages, and sex. The CD and NIBD samples are indicated by purple and red boxes. Other covariates are represented as the colors indicated by the legend.

**Supplemental Figure 2**. Colonic miRNAs separate pediatric CD into two clusters. (A) Principal component analysis (PCA) of VST normalized counts in colon tissue of pediatric CD patients (n=75) accounting for the covariates of small-RNA integrity metric (SIM), grouped ages (VEO < 6, Child = 7-12, Teen = 13-17), and sex. The two colonic clusters are represented by triangles and circles, respectively. Each sample has a color (red, green, blue, purple) according to the type of ileal disease (severe inflammation, modest inflammation, penetrating, stricturing) from each patient. The percent of variation explained is indicated for principal component 1 along the x-axis and principal component 2 along the y-axis. (B) MA plots of differentially expressed miRNAs between colon clusters 1 and 2 (baseMean > 1000, p-adj < 0.05, log2FC > 1 or -1). Dashed lines represent the log2 fold-change of expression –1.0/+1.0 (horizontal). Up- or downregulated miRNAs are colored red (up) or blue (down), with an adjusted p-val < 0.05 and baseMean > 1000. (C) A comparison of normalized read counts of three miRNAs significantly enriched in one of the colon clusters. Each data point represents a patient sample. Pink represents cluster 1 and purple represents cluster 2. (* p < 0.05, *** p< 0.001, **** p< 0.0001; Student’s t-test).

**Supplemental Figure 3**. Small intestinal architecture is unaffected by miR-29b overexpression. (A,B) Histomorphometry of 29OE/–Dox (n=4) and 29OE/+Dox (n=4) mice from initial (A) and repeated (C) experimental rounds, and of (B) wild-type (WT) mice without doxycycline treatment (WT/-Dox) (n=4) and with doxycycline treatment (WT/+Dox) (n=4). Villus height, crypt depth, and crypt density were measured for the proximal duodenum, mid-jejunum, and distal ileum. Representative brightfield H&E-stained images for villi and crypts are shown at x200 and x600 magnification, respectively. Yellow scale bars represent 100 μm at x200 magnification and 50 μm at x600 magnification. Dark blue represents the –doxycycline treatment and pink represents the +doxycycline treatment. Data presented as mean ± SEM (* p < 0.05, ** p < 0.01, *** p< 0.001, **** p< 0.0001; Student’s t-test).

**Supplemental Figure 4**. MA plot showing genes that are significantly differentially expressed from RNA-seq data in pediatric CD vs NIBD patients (n=203) in the colon (left) and ileum (right) (baseMean > 500). Dashed lines represent the log2 fold-change of expression –0.75/+0.75 (horizontal). Up- or downregulated genes are colored red or blue, respectively, with an adjusted p-val < 0.05.

**Supplemental Figure 5**. Over-expression of miR-29b reproducibly reduces the number of Paneth cells. (A) Paneth cell counts per crypt of 29OE/-Dox (n=4) and 29OE/+Dox (n=4) mice for the proximal duodenum, mid-jejunum, and distal ileum from brightfield images H&E-stained tissue sections (x600). (B) Paneth cell counts per crypt of 29OE/-Dox (n=4) and 29OE/+Dox (n=4) mice for the proximal duodenum, mid-jejunum, and distal ileum from Lyz1 immunofluorescent (red) and DAPI fluorescent (blue) images (x600). Yellow scale bars measure 50 μm. Individual Paneth cells are indicated by red arrow bars. Dark blue represents the – doxycycline treatment and pink represents the +doxycycline treatment. Data presented as mean ± SEM (* p< 0.05, ** p< 0.01,**** p< 0.0001; Student’s t-test).

**Supplemental Figure 6**. Doxycycline treatment of wild-type mice does not affect the number of Paneth cells. (A) Paneth cell counts per crypt of WT/-Dox (n=4) and WT/+Dox (n=4) mice for the proximal duodenum, mid-jejunum, and distal ileum. Representative brightfield H&E-stained images at x600 magnification. Yellow scale bar measures 50 μm. Individual Paneth cells are indicated by red arrows. Dark blue represents the –doxycycline treatment and pink represents the +doxycycline treatment. Data presented as mean ± SEM (n.s.=not significant; Student’s t-test).

**Supplemental Figure 7**. miR-29b overexpression reproducibly reduces Paneth cell marker gene expression. (A) qPCR showing the relative quantitative value (RQV) of miR-29b expression in -doxycycline (n=4) and +doxycycline (n=4) treated 29OE mice for IEC-1 and IEC-2 in a repeated experiment. Dark blue represents the –doxycycline treatment and pink represents the +doxycycline treatment. (B) qPCR of 29OE mice showing the relative quantitative value (RQV) for marker genes of intestinal stem cells (Lgr5) and four different major lineages of the intestinal epithelium (enterocyte, goblet cell, enteroendocrine cell, and Paneth cell) with and without doxycycline treatment for IEC-1 (n=4,4) and IEC-2 (n=4,4) in a repeated experiment. (C) qPCR showing the relative quantitative value (RQV) of miR-152 (C), *Copz2* (D), and *Defa17* (E) expression in -doxycycline and +doxycycline treated 29OE mice for IEC-1 (n=4,4) and IEC-2 (n=4,4) in a repeated experiment. Data presented as mean ± SEM (* p <0.05, ** p< 0.01; Student’s t-test).

**Supplemental Table 1**. Table of all clinical parameters evaluated for logistic regression analysis.

**Supplemental Table 2**. Results from the multinomial logistic regression analysis with multiple testing correction (FDR) applied to the p-values. Two clinical parameters (Rectal/sigmoid involvement and Family History of IBD) had miRNAs with significant adjusted p-values <0.1 for the colonic tissue samples. One clinical parameter (Surgery with anastomosis) had one miRNA with significant adjusted p-value< 0.1 for the ileal tissue samples.

**Supplemental Table 3**. List of 16 genes that are significantly down-regulated in both the ileum of pediatric CD patients (relative to NIBD controls) (baseMean > 150, log2FC <-0.5, padj <0.05) and in IECs of 29OE/+Dox mice (relative to 29OE/-Dox controls) (baseMean > 150, log2FC <-0.5, padj<0.05).

