## Supplemental Figures for "Aberrant miR-29 is a predictive feature of severe phenotypes in pediatric Crohn’s disease"

#### Supplemental Figure 1

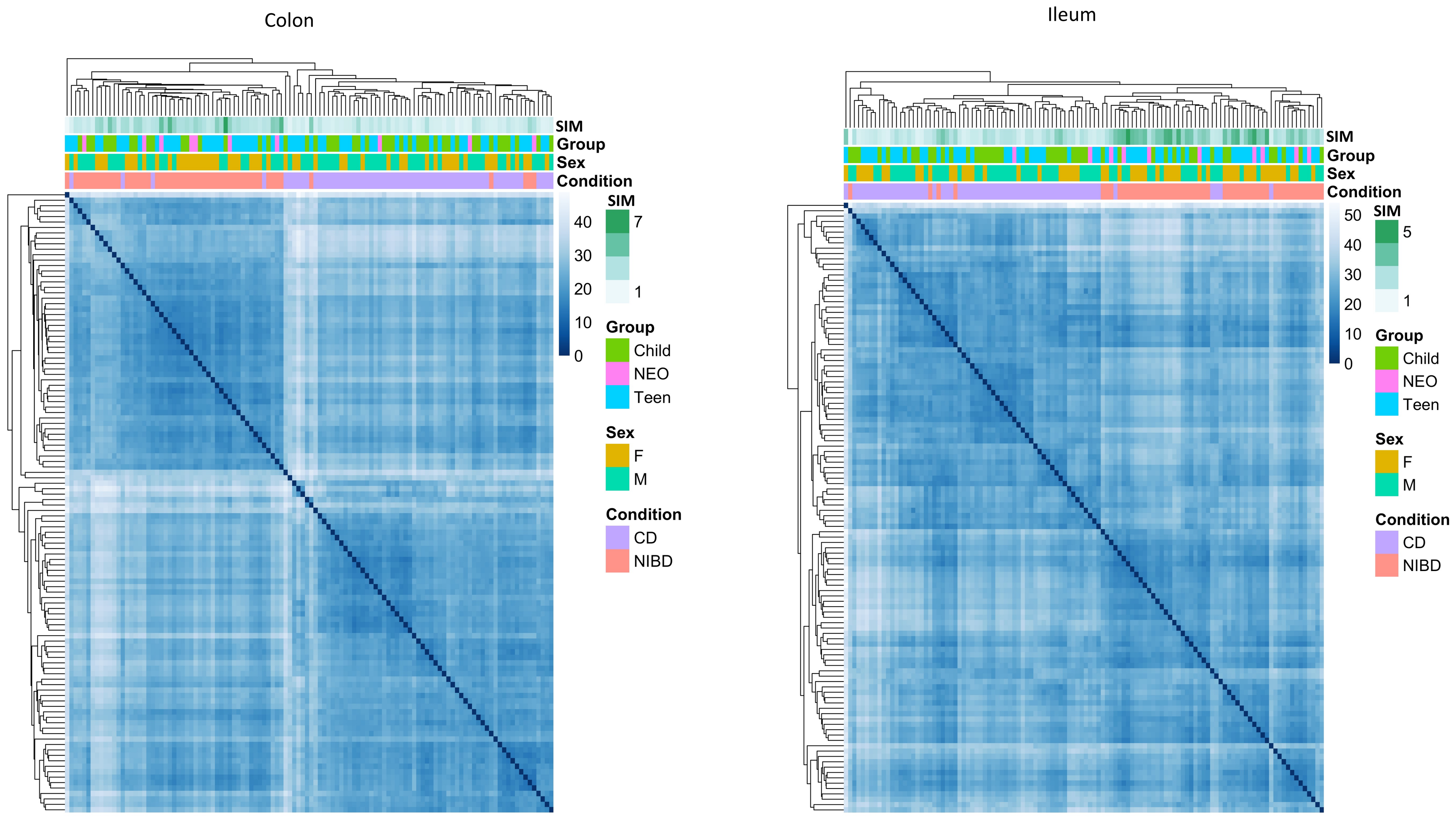

### Supplemental Figure 2

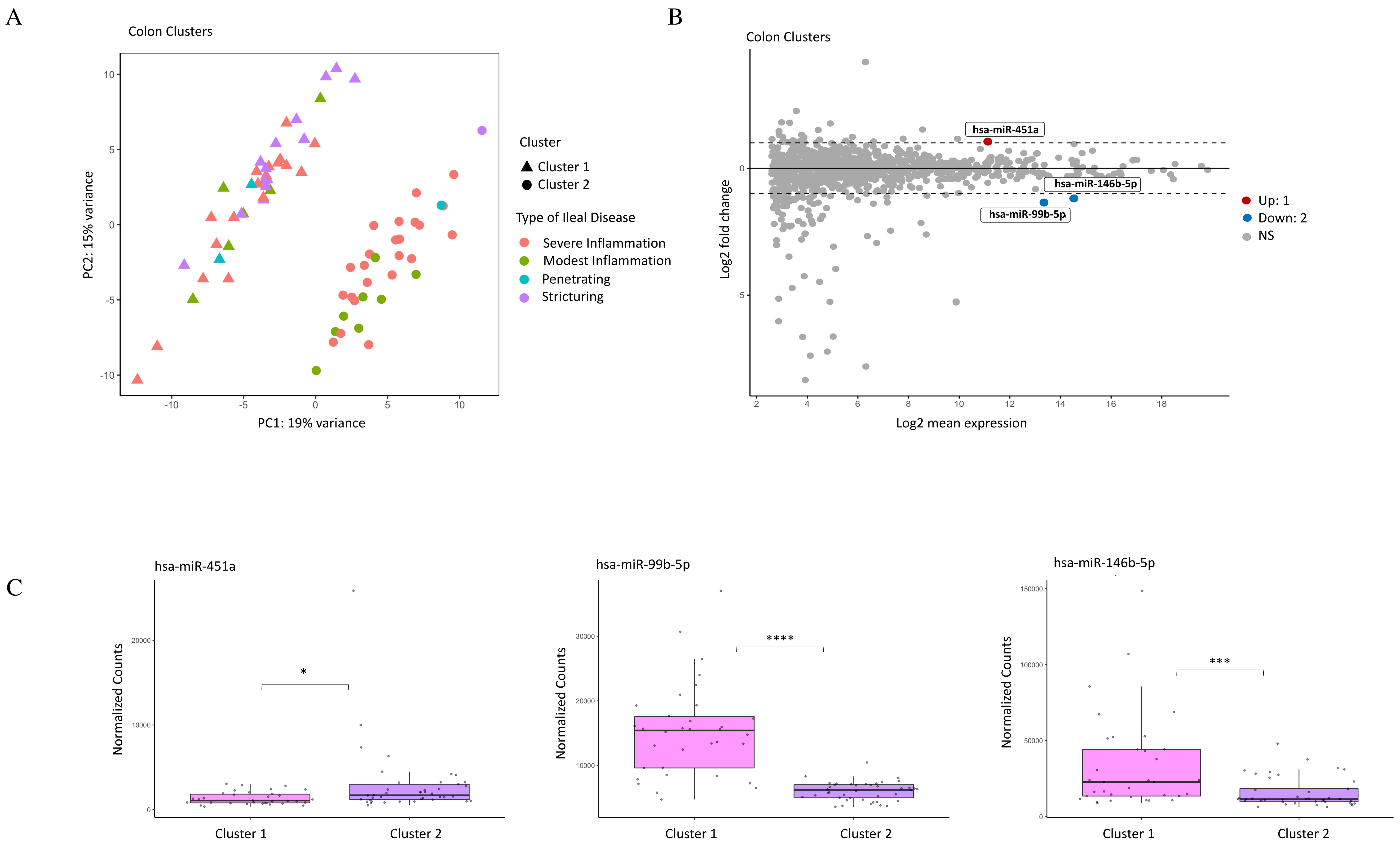

### Supplemental Figure 3

**A** 29OE First Round Histomorphometry

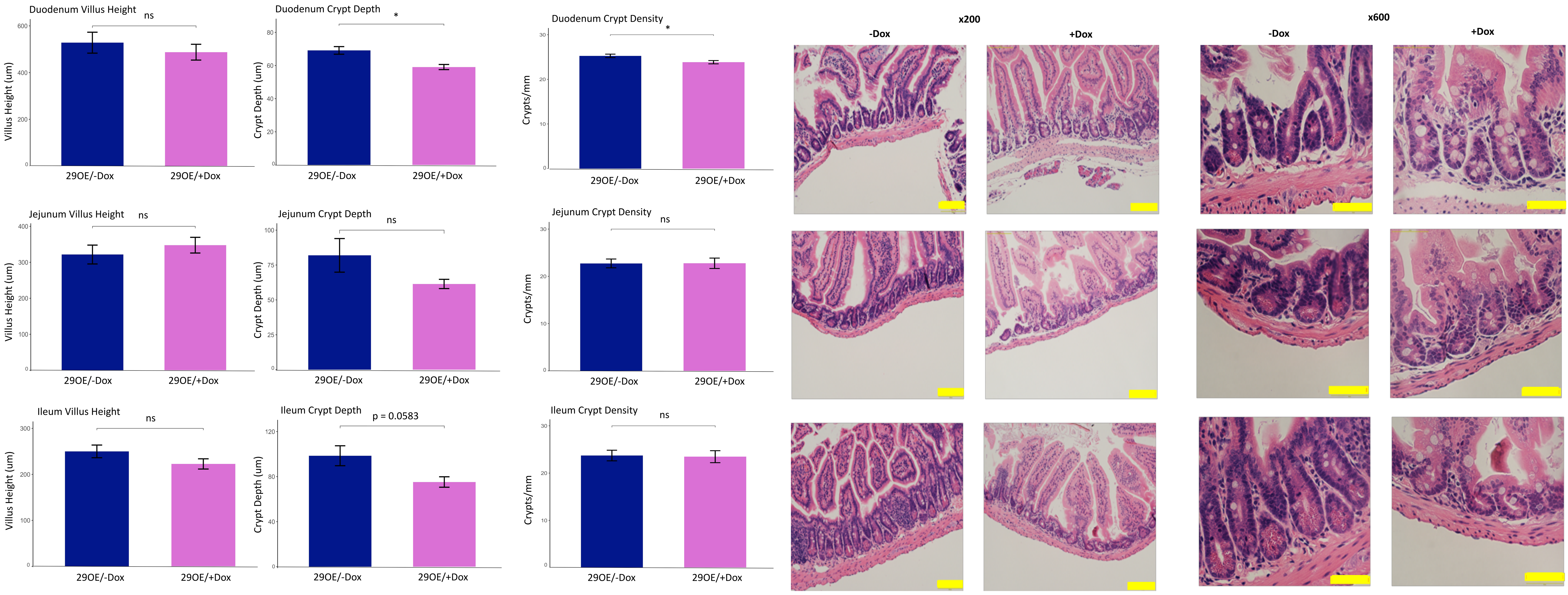


**B** WT Histomorphometry

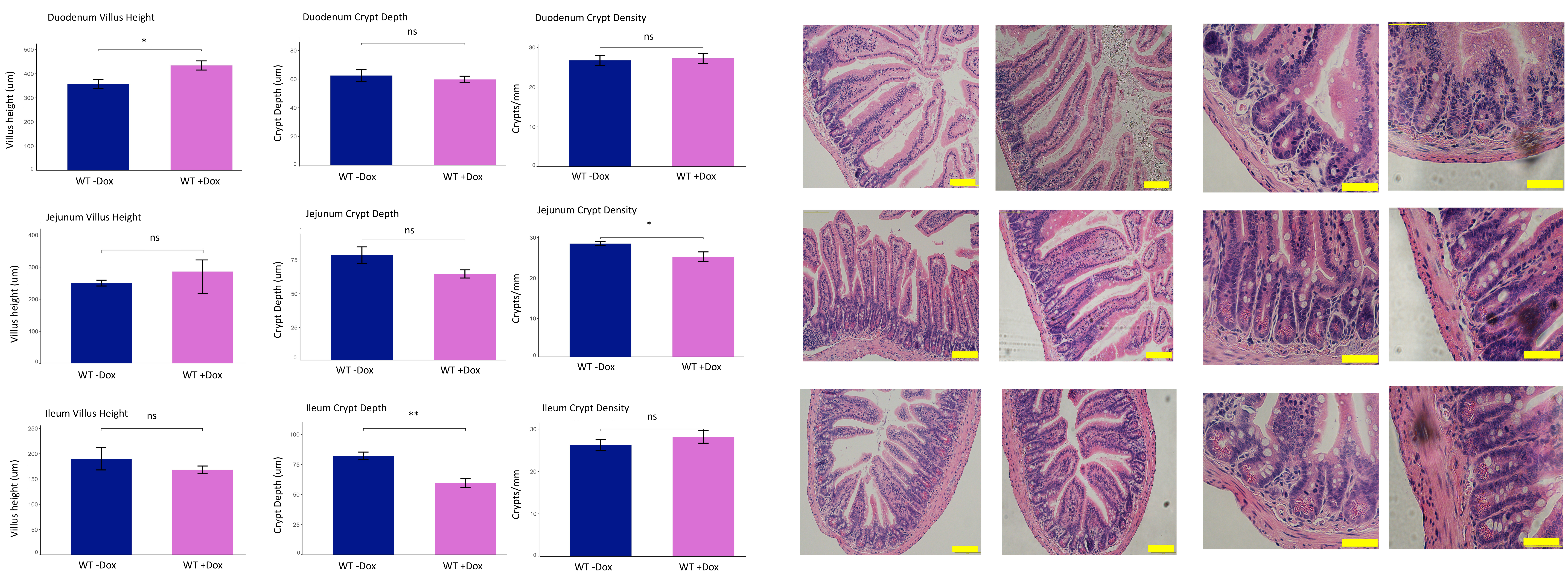

### Supplemental Figure 3 – Round 2

**C** 29OE Second Round Histomorphometry

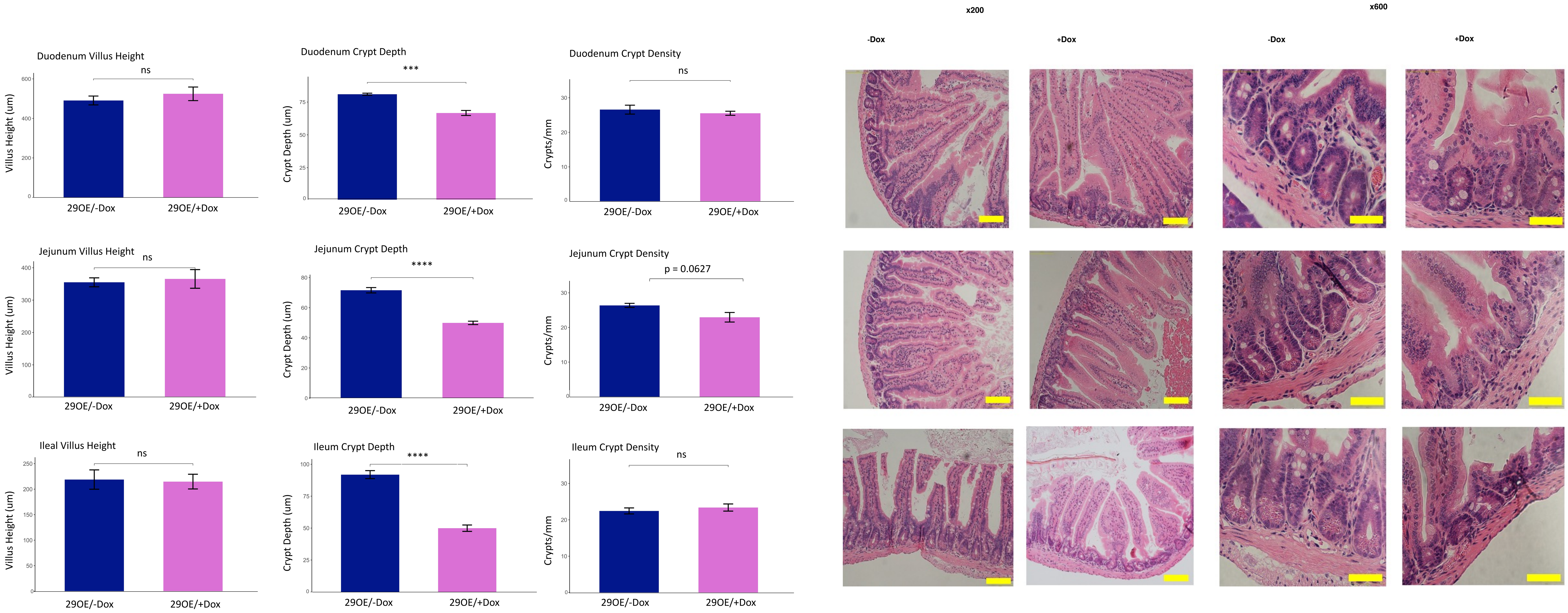

#### Supplemental Figure 4

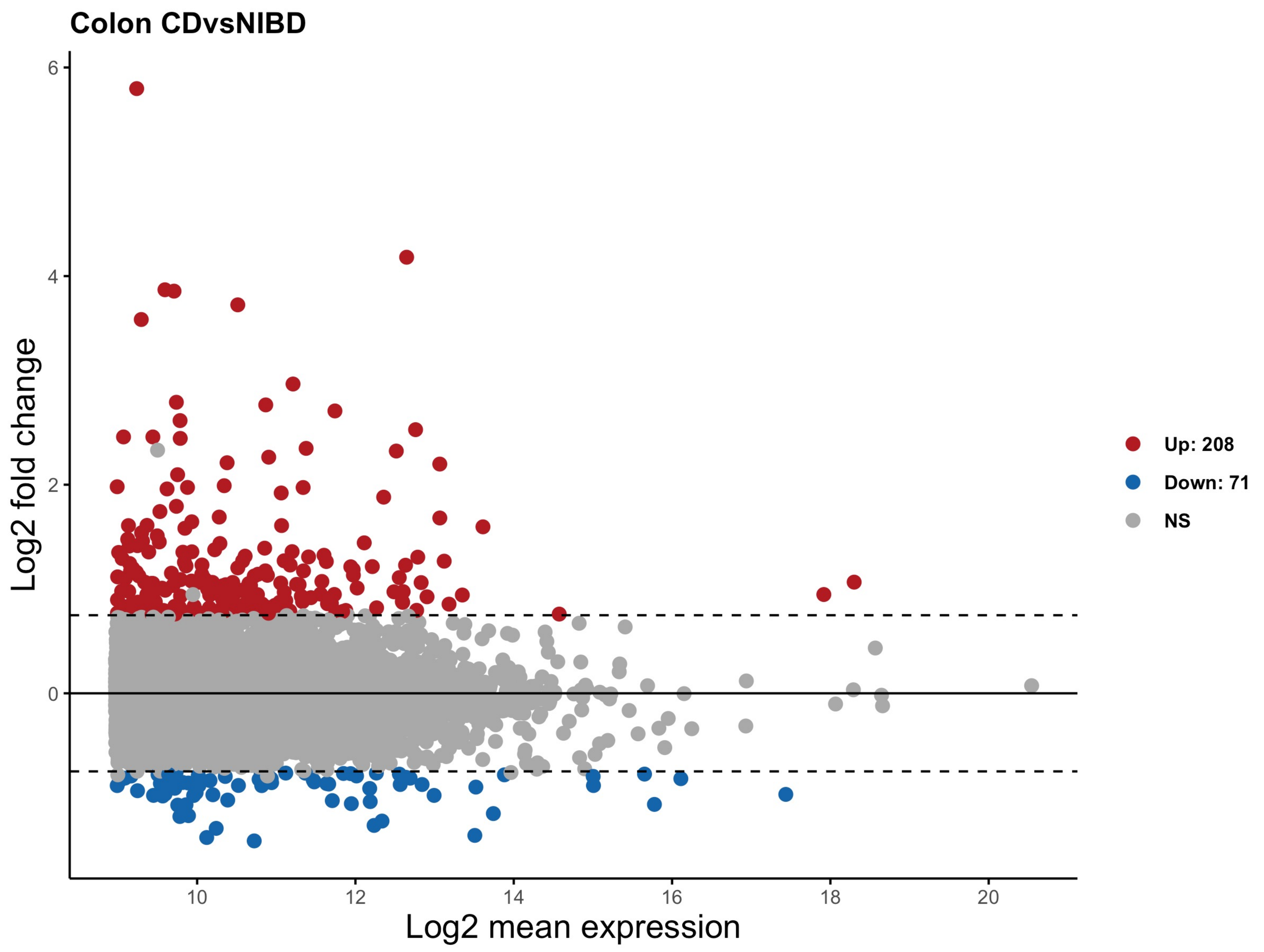


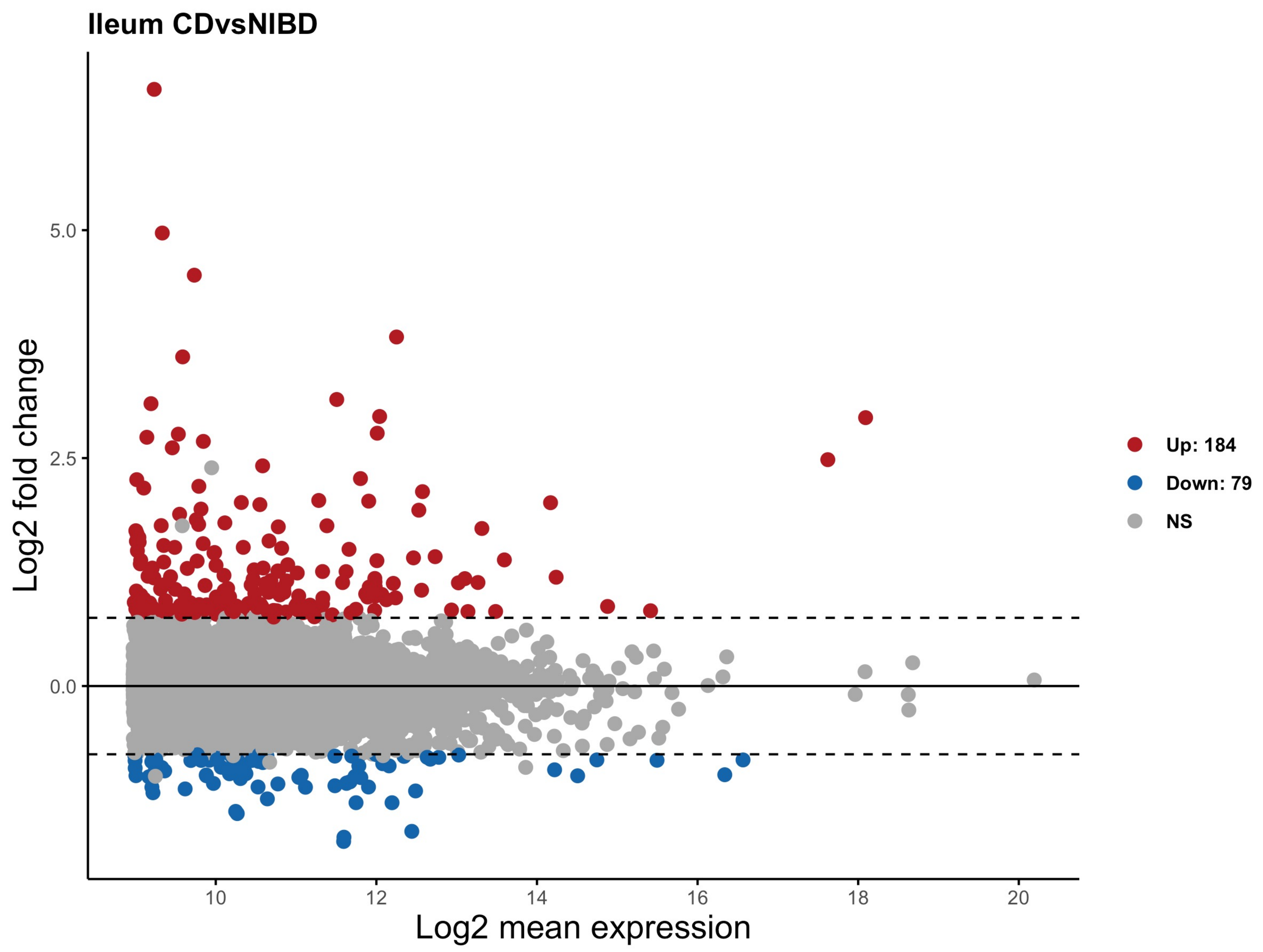

#### Supplemental Figure 5

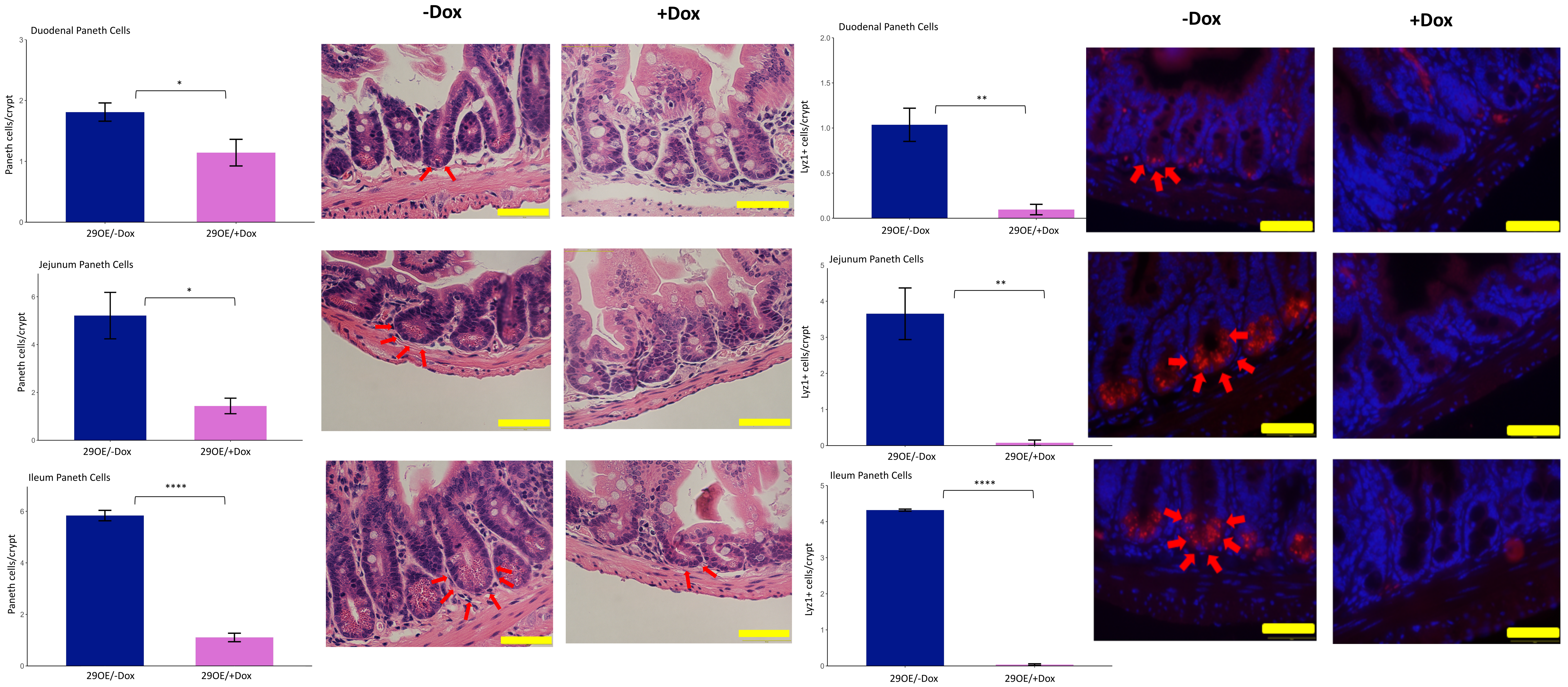

#### Supplemental Figure 6

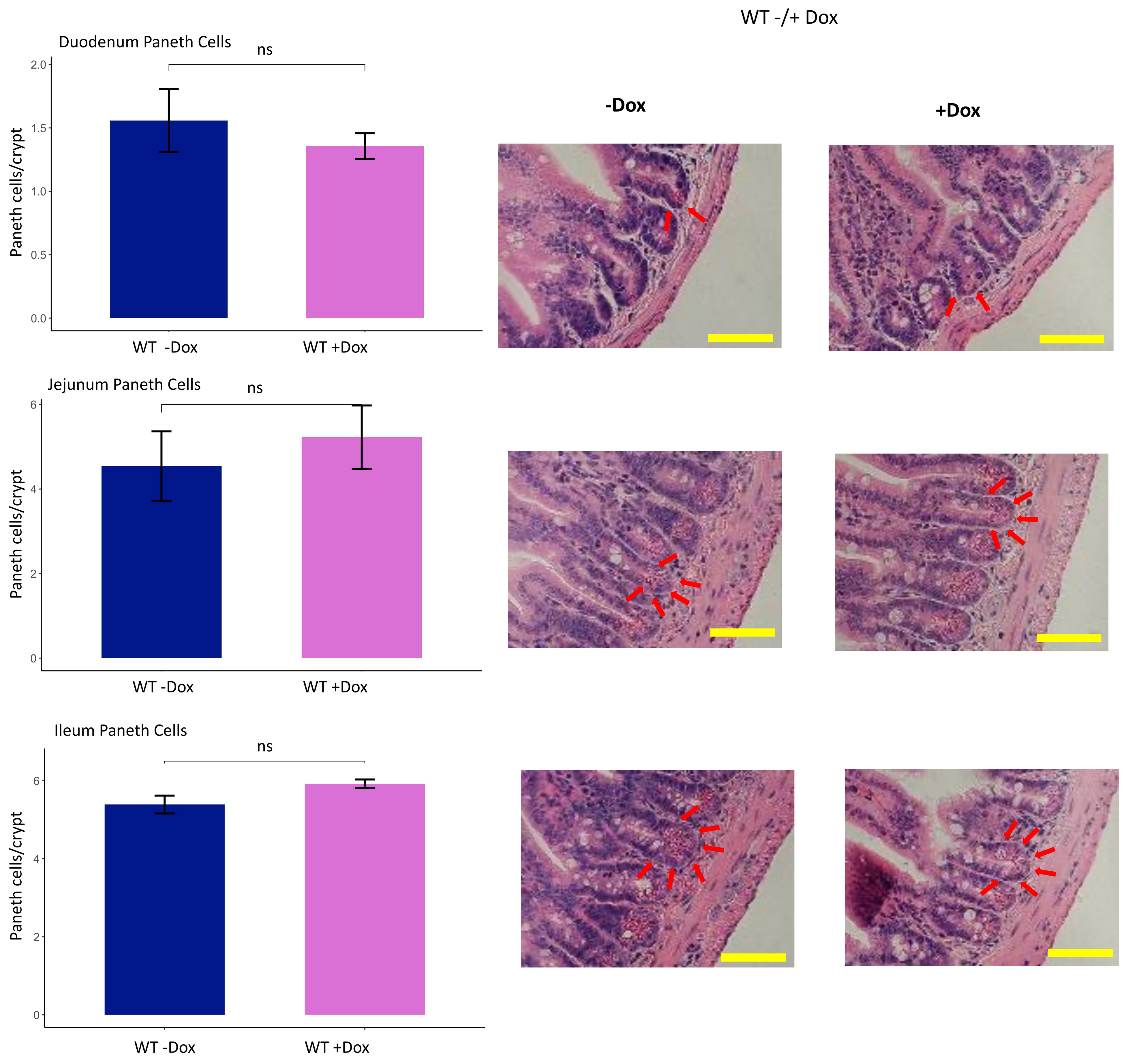

**Supplemental Figure 7**

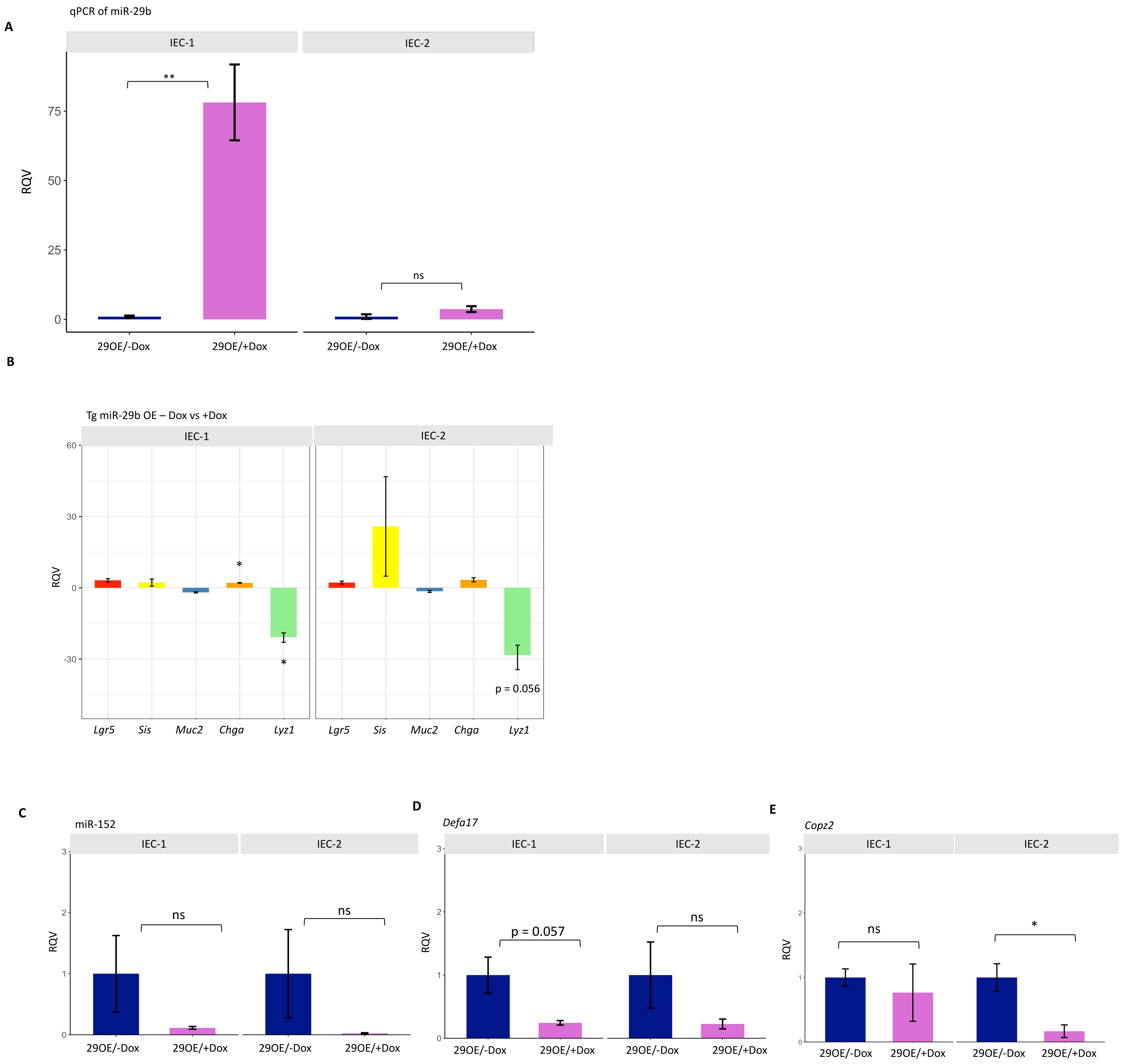

**Supplemental Table 1.** Table of all clinical parameters evaluated for logistic regression analysis.

| Clinical Parameters | Binary or Categorical | Colon CD | Ileum CD |
|---|---|---|---|
| Perianal Disease | Binary | 24 | 21 |
| Rectal or Sigmoid Involved? | Binary | 35 | 34 |
| Surgery with Anastomosis | Binary | 27 | 21 |
| Peri-anal Surgery | Binary | 11 | 10 |
| Temporary Ileostomy? | Binary | 6 | 5 |
| # of times therapy escalated (Average) | Categorical | 1.32 | 1.27 |
| Current Biologic? | Binary | 40 | 33 |
| Current IM? | Binary | 39 | 35 |
| Remission? | Binary | 58 | 50 |
| Sex | Binary | F = 27<br>M = 48 | F = 22<br>M = 43 |
| Diagnosis Age | Categorical | VEO = 2<br>Child = 36<br>Teen = 37 | VEO = 3<br>Child = 31<br>Teen = 31 |
| Family History of IBD | Binary | 30 | 25 |
| Locations | Categorical | L1 = 11<br>L2 = 11<br>L3 = 49<br>L4 = 4 | L1 = 10<br>L2 = 8<br>L3 = 43<br>L4 = 4 |
| Failed IM? | Binary | 31 | 26 |
| Type of Ileal Disease | Categorical | Modest Inflammation =15<br>Severe Inflammation = 41<br>Stricturing = 16<br>Penetrating = 3 | Modest Inflammation =13<br>Severe Inflammation = 38<br>Stricturing =13<br>Penetrating = 1 |

**Supplemental Table 2** Results from the multinomial logistic regression analysis with multiple testing correction (FDR) applied to the p-values. Two clinical parameters (Rectal/sigmoid involvement and Family History of IBD) had significant adjusted p-values with p< 0.1 for the colonic tissue samples. One clinical parameter (Surgery with anastomosis) had one microRNA with significant adjusted p-values with p< 0.1 for the ileal tissue samples

| Colonic microRNA | Rectal or Sigmoid Involved |
|---|---|
| hsa-miR-21-5p | 0.00508425 |
| hsa-miR-21-5p_+_1 | 0.00508425 |
| hsa-miR-21-3p | 0.01109579 |
| hsa-miR-31-5p | 0.01416445 |
| hsa-miR-26a-5p_-_1 | 0.05250646 |
| hsa-let-7b-5p | 0.07157595 |
| hsa-miR-215_-_1 | 0.07157595 |

| Colonic microRNA | Family History |
|---|---|
| hsa-miR-142-5p | 0.0703179 |
| hsa-miR-142-5p_+_1 | 0.0703179 |
| hsa-miR-16-5p | 0.0703179 |
| hsa-miR-215_-_1 | 0.0703179 |
| hsa-miR-29a-3p | 0.0703179 |
| hsa-miR-15a-5p | 0.07101881 |
| hsa-miR-142-3p_+_3 | 0.08506924 |
| hsa-miR-451a | 0.08506924 |

| Ileal microRNA | Surgery with Anastomosis |
|---|---|
| hsa-miR-215_-_1 | 0.09907668 |

**Supplemental Table 3.** List of 16 genes that are significantly down-regulated in both the ileum of pediatric CD patients (relative to NIBD controls) (baseMean > 150, log2FC <-0.5, padj <0.05) and in IECs of 29OE/+Dox mice (relative to 29OE/-Dox controls) (baseMean > 150 , log2FC <-0.5 , padj<0.05).

| Gene | Species/Tissue Located in |
|---|---|
| DSC2 | Pediatric Colon, Pediatric Ileum, OE 29 Mouse |
| EPB41L4B | Pediatric Colon, Pediatric Ileum, OE 29 Mouse |
| EPCAM | Pediatric Colon, Pediatric Ileum, OE 29 Mouse |
| HSPA1B | Pediatric Colon, Pediatric Ileum, OE 29 Mouse |
| LGALS3 | Pediatric Colon, Pediatric Ileum, OE 29 Mouse |
| PRR15 | Pediatric Colon, Pediatric Ileum, OE 29 Mouse |
| ACSF2 | Pediatric Ileum, OE 29 Mouse |
| DNASE1 | Pediatric Ileum, OE 29 Mouse |
| ENPP3 | Pediatric Ileum, OE 29 Mouse |
| FAM151A | Pediatric Ileum, OE 29 Mouse |
| NR1D2 | Pediatric Ileum, OE 29 Mouse |
| PDZK1 | Pediatric Ileum, OE 29 Mouse |
| PMP22 | Pediatric Ileum, OE 29 Mouse |
| SLC52A3 | Pediatric Ileum, OE 29 Mouse |
| SLC5A1 | Pediatric Ileum, OE 29 Mouse |
| XPNPEP2 | Pediatric Ileum, OE 29 Mouse |
